# Repurposed macroH2A Chromatin Landscapes Guide Cellular Reprogramming

**DOI:** 10.64898/2025.12.11.693653

**Authors:** Dimitrios Valakos, Antonis Kokkalis, Eleftheria Klagkou, Alexander Polyzos, Giannis Vatsellas, Dimitris Thanos

## Abstract

Cellular reprogramming converts differentiated cells into a pluripotent state through extensive chromatin remodeling. The histone variant macroH2A has classically viewed as an epigenetic barrier stabilizing somatic identity and restricting pluripotency gene activation. Here, we show that during reprogramming, macroH2A1 nucleosomes undergo functional repurposing. Early in the process, mH2A1 nucleosomes act as a barrier to cellular plasticity, but later facilitate the establishment and maintenance of pluripotency by reshaping the epigenetic landscape. High-resolution chromatin profiling reveals that mH2A1.2 nucleosomes undergo rapid, large-scale repositioning, dissociating from promoters and reassembling ∼30 bp away, frequently near NRF-1 binding sites. This repositioning occludes E2F4 binding, relieving cell-cycle arrest thereby enabling reprogramming. Likewise, mH2A1.1 nucleosomes display extensive mobility that culminates in deposition at pluripotency genes in ESCs, where they sustain their expression by assembling promoter transcriptional hubs. In this later role, mH2A1.1 functions as a chromatin bookmark stabilizing the Nanog, Sox2, and Oct4 network. These findings suggest that mH2A1 nucleosomal mobility underlines its context-dependent functional repurposing from reprogramming inhibitor to facilitator, illustrating how chromatin components evolve dynamic roles to coordinate cell-state transitions.

## Introduction

Gene expression programs are dynamically regulated during development, throughout adult life, and in response to environmental cues. The molecular basis of these complex regulatory networks lies in the interplay between cis-acting regulatory elements and transcription factors (TFs), which mediate the recruitment of additional transcriptional regulatory proteins such as cofactors, chromatin modifiers, and the transcriptional machinery at gene promoters in a cell type-specific manner within specialized chromatin landscapes (Merika & Thanos, 2001; Chen & Dent, 2014; Reiter *et al*, 2017; Agelopoulos & Thanos, 2006; Donohue *et al*, 2022; Kim & Wysocka, 2023; Park *et al*, 2025). A critical transcriptional control layer is exerted by nucleosomes on regulatory elements whose exact positioning, disassembly/reassembly, or sliding can determine transcriptional activity by modulating the accessibility of regulatory DNA sequences to TFs (Whitehouse *et al*, 1999; Lomvardas & Thanos, 2001; Kassabov *et al*, 2003; Yuan *et al*, 2005; Brogaard *et al*, 2012; Apostolou & Hochedlinger, 2013; Soufi *et al*, 2015; Nocetti & Whitehouse, 2016; Rudnizky *et al*, 2019; Harwood *et al*, 2019; Oruba *et al*, 2020; Gamarra & Narlikar, 2021; Mohammed Ismail *et al*, 2023; Grand *et al*, 2024; Duttke *et al*, 2024).

Histone variants, particularly those of H2A, contribute to chromatin complexity by forming specialized nucleosomes at specific genomic locations throughout the cell cycle, thus conferring distinct biochemical properties with unique biological consequences (Turinetto & Giachino, 2015). The histone variant macroH2A (mH2A) consists of an H2A-like region, followed by a linker and a large globular macro domain (Pehrson & Fried, 1992). This variant is highly conserved and widespread in vertebrates, encoded by two distinct genes in mammals, *H2afy* and *H2afy2*. *H2afy* produces the mH2A1 protein, which exists in two splicing isoforms, mH2A1.1 and mH2A1.2. Notably, only mH2A1.1 can bind ADP- and polyADP-ribose (Karras *et al*, 2005), linking it to metabolic processes, DNA damage response pathways, and apoptosis (Timinszky *et al*, 2009; Kozlowski *et al*, 2018). *H2afy2* encodes the mH2A2 variant (Costanzi & Pehrson, 2001).

Although mH2A was first linked to gene repression due to its enrichment on the inactive X chromosome, it is now known to be broadly distributed across heterochromatin and euchromatin (Costanzi & Pehrson, 2001; Agelopoulos & Thanos, 2006; Schones *et al*, 2008; Teif *et al*, 2012; Carone *et al*, 2014; Lavigne *et al*, 2015; Sun *et al*, 2018; Sun & Bernstein, 2019). mH2A alters nucleosome architecture and strongly inhibits TF binding to their DNA recognition sites. When mH2A-nucleosomes mask activator binding sites, they typically lead to gene repression, whereas masking of repressor sites can result in transcriptional derepression (Lavigne *et al*, 2015; Ni *et al*, 2020). Through these context-dependent effects, mH2A reduces transcriptional noise and stabilizes gene expression programs (Lavigne *et al*, 2015), helping to preserve cell identity (Valakos *et al*, 2023). Knockdown studies showed that mH2A1 and mH2A2 act as barriers to reprogramming (Pasque *et al*, 2011; Gaspar-Maia *et al*, 2013; Barrero *et al*, 2013), largely by blocking Mesenchymal to Epithelial Transition (MET) and maintaining mesenchymal gene expression networks, including extracellular matrix, cytoskeletal, membrane, and regulatory genes (Pliatska *et al*, 2018; Valakos *et al*, 2023). Still, these findings do not rule out additional roles for mH2A variants at later stages of reprogramming, which have remained elusive because of their dominant early effects.

Beyond reprogramming, mH2A variants influence Epithelial to Mesenchymal Transition (EMT) in cancer metastasis (Hodge *et al*, 2018) and function as tumor suppressors in melanoma, lung, and breast cancers (Sporn *et al*, 2009; Kapoor *et al*, 2010; Xu *et al*, 2015; Broggi *et al*, 2020). They downregulate Cdk8 and suppress E2F1, thereby restraining cell-cycle progression (Buschbeck *et al*, 2009; Kapoor *et al*, 2010;)—an axis central to both tumorigenesis and induced Pluripotent Stem Cells (iPSC) generation. These findings highlight mH2A as a critical regulator of cell-cycle progression—a process essential for both cancer formation and the generation of induced pluripotent stem cells (iPSCs).

To clarify the molecular functions of mH2A during the entire course of reprogramming, we performed time-resolved ChIP-seq profiling. We uncovered a striking and unprecedented genome-wide redistribution of mH2A1—but not mH2A2—nucleosomes, pointing to a previously unknown function of mH2A1 in shaping dynamic time-dependent epigenetic landscapes. Promoter-bound mH2A1.2 nucleosomes in MEFs rapidly dissociate at the onset of reprogramming and later reappear at nearby but shifted positions (∼30 bp), particularly on cell-cycle gene promoters. This repositioning correlates with NRF-1 binding and results in the masking of E2F4 binding motifs. In MEFs, E2F4 binds these promoters to enforce cell-cycle arrest and its access is permitted by the original nucleosome position. However, in ESCs/iPSCs, that represent the end point of reprogramming, the slight positional shift of mH2A1.2 nucleosomes blocks E2F4 from accessing its binding sites, releasing the arrest and enabling proliferation. In parallel, we uncovered a novel function for mH2A1.1 as a bookmark for the maintenance of the pluripotency network in embryonic stem cells (ESCs). Overall, these findings reveal how dynamic mH2A1 nucleosome mobility repurposes its biological function to regulate cell-cycle control and pluripotency during reprogramming, a discovery with broader implications in stem cell biology and cancer.

## Results

### Genomic Co-Occupancy of mH2A1.1 Nucleosomes and OSKM in ESCs

Since mH2A-nucleosomes can establish a specialized chromatin environment acting as a barrier to somatic cellular reprogramming, we sought to determine whether mH2A-bound DNA-regions overlap with Oct4, Sox2, Klf4 and c-Myc (OSKM)-binding sites, and thus prevent their DNA binding. To test this hypothesis, we systematically analyzed the genomic regions occupied by mH2A1.1-, mH2A1.2-, and mH2A2-nucleosomes in comparison to OSKM-binding during the course of cellular reprogramming. We performed ChIP-seq experiments using native chromatin digested with micrococcal nuclease (MNase) from MEFs undergoing reprogramming at Days 0, 1, 2, 3, 6, and 9, all of which represent intermediate stages of reprogramming, as well as from Bruce-4 ESCs corresponding to the end point of reprogramming. We have previously shown the close resemblance between the transcriptome profile of Bruce-4 ESCs and that of iPSCs generated in our reprogramming experiments (Papathanasiou *et al*, 2021). This approach allowed us to identify at near base-pair resolution the position of mH2A-nucleosomes at each time-point of reprogramming and compare their relative location between different time-points of reprogramming (Schones *et al*, 2008; Brogaard *et al*, 2012; Teif *et al*, 2012; Carone *et al*, 2014; Nocetti & Whitehouse, 2016; Voong *et al*, 2016; Oruba *et al*, 2020; Piroeva *et al*, 2023). Immunoprecipitation was carried out with our mH2A variant-specific antibodies (α-mH2A1.1, α-mH2A1.2, and α-mH2A2) (Valakos *et al*, 2023), followed by deep DNA sequencing and extensive computational analysis. Control experiments demonstrating the specificity of our antibodies have been previously described (Valakos *et al*, 2023). Our goal was to obtain a high-resolution comparative genome-wide map between mH2A-nucleosomes and OSKM binding, offering new insights into the role of specialized chromatin landscapes in transcription factor accessibility during cellular reprogramming.

We profiled and compared the *in vivo* binding of individual OSKM factors (Klagkou *et al*, 2024) to mH2A1.1 and mH2A1.2 nucleosome-bound regions and surprisingly we found a strong and highly specific co-occupancy between mH2A1.1-nucleosomes and OSKM in ESCs, with overlap percentages ranging from 36.41% (Oct4) to 48.67% (Myc) (Fig. 1A, left panel). In contrast, no significant overlap was detected during the intermediate stages of reprogramming (Days 1, 2, 3, 6, and 9). Interestingly, prior to reprogramming (MEFs/Day 0), Myc and to a lesser extent, Klf4 binding events were co-localized with mH2A1.1-nucleosomes, but this association was reduced or lost immediately upon initiation of reprogramming (Day 1), and re-emerged after the establishment of pluripotency in ESCs (Fig. 1A, left panel). These findings indicate a dynamic and stage-specific interplay between mH2A1.1-nucleosomes and KM-binding at the beginning and the end of reprogramming (Fig. 1A, left panel). By contrast to mH2A1.1, mH2A1.2-nucleosomes do not substantially overlap with OSKM in ESCs (Fig. 1A, right panel). These data suggest that during reprogramming, OSKM factors bind to the genome largely independent of mH2A1.1- and mH2A1.2-nucleosomes, but upon the establishment of pluripotency, approximately 50% of the total OSKM-binding events occur predominantly to mH2A1.1-nucleosomal-bound genomic regions. This mH2A1.1 nucleosome-binding property of OSKM aligns with their pioneer activity since they can bind to their DNA motifs within nucleosomes (Soufi *et al*, 2015). Taken together, our data reveal an unprecedented interplay linking mH2A1.1-nucleosome positioning to OSKM-binding in ESCs, suggesting a previously unsuspected regulatory role in pluripotent cells.

**Figure 1:**
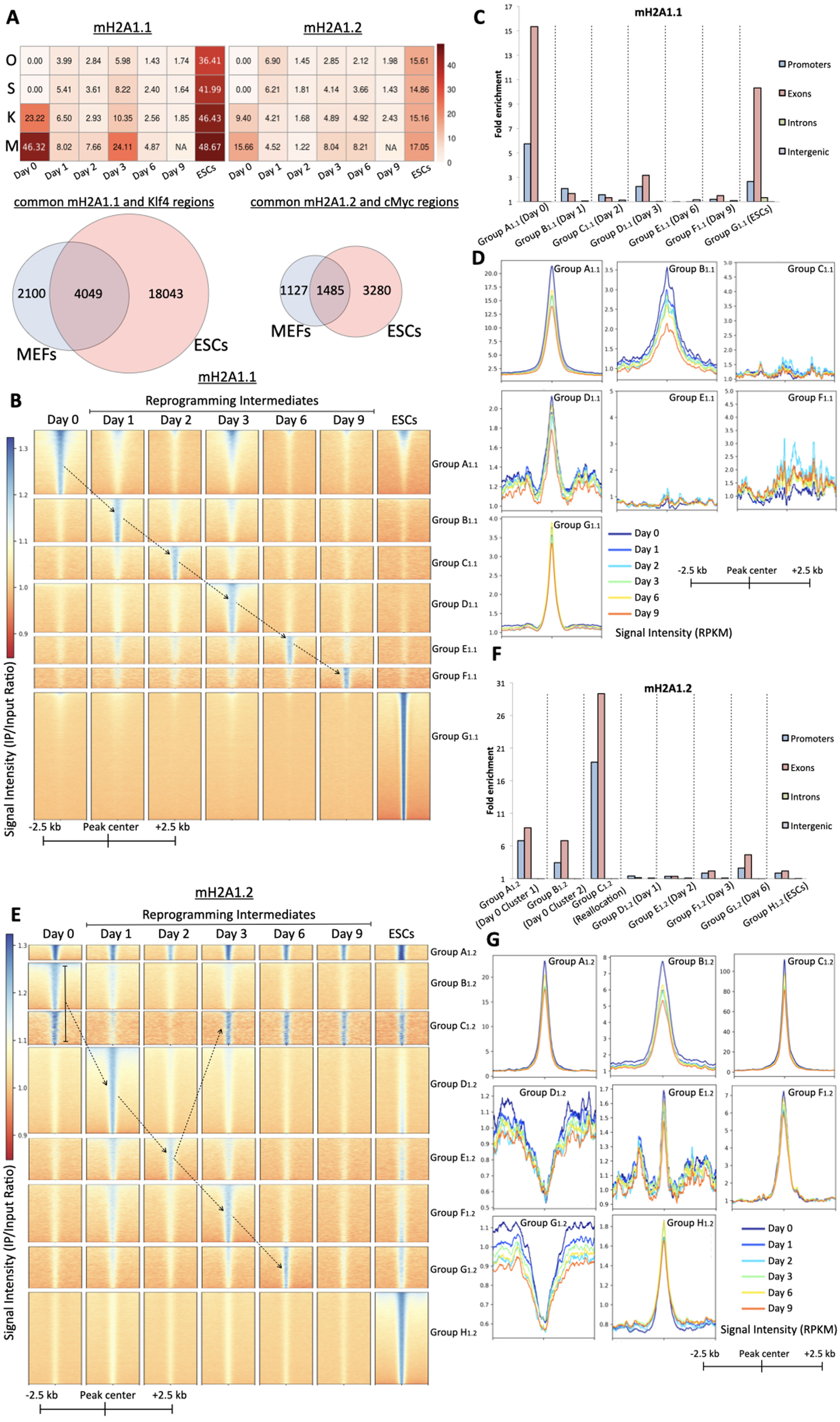
Dynamic reconfiguration of the mH2A1.1 and mH2A1.2 chromatin landscapes during cellular reprogramming. (A) Top: Heatmaps showing the percentage of shared binding peaks between each of the OSKM factors (listed on the left) and either mH2A1.1 (left) or mH2A1.2 (right) nucleosomes at multiple stages of reprogramming (Day 0, Day 1, Day 2, Day 3, Day 6, Day 9) and ESCs. Bottom: Venn diagrams illustrating the overlap between genomic regions bound by mH2A1.1 and Klf4 (left) or mH2A1.1 and c-Myc (right) in MEFs and ESCs, respectively. NA: Not Available (B) Heatmaps displaying the genome-wide redistribution of mH2A1.1-nucleosomes across distinct genomic regions (Groups A_1.1_–G_1.1_), from –2.5 kb to +2.5 kb relative to peak centers, throughout reprogramming (Day 0 to ESCs). Regions are grouped based on dynamic mH2A1.1 occupancy patterns. Peaks were called using MACS2 and signal intensities are shown as IP/Input ratio. Dashed arrows represent the dynamic relocation of mH2A1.1 nucleosomes between time-points. (C) Bar graph showing the fold enrichment in genomic annotations of mH2A1.1 Groups A_1.1_–G_1.1_ regions (as defined in Fig. 1B), categorized into promoters, exons, introns and intergenic regions. (D) Line plots summarizing ATAC-seq signal (RPKM) across Groups A_1.1_–G_1.1_ (defined in B) from –2.5 kb to +2.5 kb relative to mH2A1.1 peak centers, shown for MEFs (Day 0), Day 1, Day 2, Day 3, Day 6 and Day 9 of reprogramming. (E) Same as (B), but for mH2A1.2 nucleosomes. (F) Same as (C), showing the fold enrichment in genomic annotations within the mH2A1.2 occupancy groups defined in (E). (G) Same as (D), depicting ATAC-seq signal across mH2A1.2 groups from (E).

### Dynamic redistribution of mH2A1.1 and mH2A1.2 nucleosomes during reprogramming

Since the mH2A1 chromatin landscape is not generally linked to OSKM DNA-binding during the intermediate stages of reprogramming, we tracked the fate of mH2A1-nucleosomes positioning throughout the process and intersected these data with TF binding data and transcriptomics changes. This was motivated mainly by the fact that mH2A-nucleosomes generally function as inhibitors of reprogramming by blocking MET (Pliatska *et al*, 2018). This inhibitory effect might be the result of mH2A-nucleosomes acting either as barriers to TF binding, or inhibiting chromatin remodeling by altering nucleosome structure, or recruiting repressive factors, or by a combination thereof. Strikingly, we discovered that the mH2A1-nucleosome landscape is marked by an unprecedented fluidity characterized by highly dynamic patterns of sequential genomic occupancies and ejections en route to pluripotency (Fig. 1B and 1E, Datasets EV1 and EV2). More specifically, we found that most of the genomic sites that are strongly occupied by mH2A1.1–nucleosomes in naïve MEFs (Day 0, Group A_1.1_) eject these nucleosomes on the first day of reprogramming, followed by association to a novel set of sites (Fig. 1B, compare Group A_1.1_ and Group B_1.1_ at Days 0 and 1). Again, on Day 2, most of the newly occupied Group B_1.1_ nucleosomes depart and reappear to a novel set of genomic sites (Group C_1.1_), departing again from them and appearing elsewhere on Day 3 (Group D_1.1_) and a fraction of them reappearing on Group A_1.1_. Importantly, on Days 6 and 9, the fraction of the reprogrammable genome occupied by mH2A1.1 nucleosomes is reduced by ∼50% of that in MEFs (Fig. 1B, compare height of Group A_1.1_ to height of Groups E_1.1_ and F_1.1_). However, in ESCs, which represent the end point of reprogramming, we found that nearly all mH2A1.1-nucleosomes occupy a novel and significantly larger set of genomic sites that were not previously associated with mH2A1.1 nucleosomes (Group G_1.1_). Notably, a very small number of MEF mH2A1.1-nucleosomes reappear in ESCs (Fig. 1B, Group A_1.1_). Taken together, our findings reveal that most of the mH2A1.1-nucleosomes in MEFs depart almost immediately from their original location at the beginning of reprogramming and are gradually relocated to new sites until they reach their final destination in terminal pluripotent cells.

Next, we assigned the highly mobile mH2A1.1-nucleosomes relative to genomic features and found that Groups A_1.1_ and G_1.1_, which are mostly found in MEFs and ESCs, respectively, are enriched in promoter regions (Fig. 1C, Fig. EV1A). This implies that a fraction of the mH2A1.1-nucleosomes occupying promoters in MEFs depart and land on a different set of promoters in ESCs, and thus could influence the expression of the corresponding genes. The Group A_1.1_ genomic sites (found in both MEFs and many of those preserved in ESCs) are related to genes with house-keeping functions like mRNA processing and translation (Fig. EV1B, Group A_1.1_ panel), whereas Group G_1.1_ (ESCs) associated genes are directly related to pluripotency mechanisms (Fig. EV1B, lower-right panel), a finding suggesting a role for mH2A1.1 in maintenance of pluripotency. ATAC-seq analysis verified that the Groups A_1.1_ and G_1.1_ are highly accessible (Fig. 1D and Fig. EV1C), while monitoring the Group G_1.1_ sites (ESCs) during reprogramming revealed that many of them preexist in an open configuration in MEFs (Fig. 1D).

Similar to mH2A1.1-nucleosomes, we uncovered a dynamic but distinct pattern of mH2A1.2-nucleosomes relocation during cellular reprogramming (Fig. 1E). We identified three separate groups of regions bound by mH2A1.2 in MEFs (Group A_1.2,_ Group B_1.2_ and Group C_1.2,_ Fig. 1E). Although Group A_1.2_ regions are stably bound by mH2A1.2 nucleosomes throughout reprogramming, most nucleosomes in Groups B_1.2_ and C_1.2_ depart at the beginning of reprogramming (Day 1) and are either permanently destabilized (Group B_1.2_) or reappear later (Group C_1.2_) (Fig. 1E). Similar to mH2A1.1, various regions of the genome are sequentially occupied by mH2A1.2-nucleosomes during reprogramming. An interesting difference between mH2A1.1 and mHA1.2 dynamic nucleosome relocation patterns is that the former are changing their locations more abruptly during reprogramming, as opposed to mH2A1.2-nucleosomes, whose binding, with the exception of Group C_1.2_, appears less distinguishable between time-points (compare the panels with the different time-points in Fig. 1B and 1E). These observations suggest that the molecular mechanisms responsible for mH2A1 chromatin landscape fluidity can discriminate between the mH2A1.1 and mH2A1.2 nucleosomes.

Specifically, Group C_1.2_ regions depict strong, sharp binding on Day 0, which resolves on Days 1 and 2 and reappears on Day 3, remaining stable until pluripotency (Fig. 1E). Group C_1.2_ regions are highly enriched in promoter elements (Fig. 1F, Fig. EV2A), having the highest promoter enrichment and the highest chromatin accessibility in ESCs between all Groups of both mH2A1.1 and mH2A1.2 datasets (compare Fig. 1C with Fig. 1F, Fig. 1D with Fig. 1G and Fig. EV1C with Fig. EV2C). Group C_1.2_-related genes are involved in metabolism, protein transport and cell-cycle pathways (Fig. EV2B).

In sharp contrast to the highly mobile mH2A1-nucleosomes during reprogramming, the mH2A2-nucleosomes are significantly less mobile, as genomic sites bound by mH2A2-nucleosomes in MEFs are maintained during the first 6 days of reprogramming (Fig. EV3A Group A_2_, Dataset EV3), while a new group of sites (Group B_2_ region) are progressively occupied during reprogramming reaching maximum mH2A2-binding in ESCs. Furthermore, mH2A2-nucleosomes do not share strong overlaps with the OSKM-binding sites (Fig. EV3B), they build largely inaccessible chromatin structures (Fig. EV3C) and are not enriched in promoters (Fig. EV3D).

So far, our results indicate that the mH2A1 chromatin landscape is remarkably flexible, a property inconsistent with mH2A1-nucleosomes inhibitory effect on reprogramming. Notably, comparison of the genomic sites occupied by the continuously flowing mH2A1.1- and mH2A1.2-nucleosomes revealed that they share a relatively small number of common sites (Fig. EV2D), thus further suggesting the involvement of chromatin remodeling machines that can distinguish between the two isoforms. In fact, the actual number of the common sites is overestimated due to the fact that mH2A ChIP-seq peak-calling was performed using broad peak-calling modes, which is the optimal method to be used for histones and histone modifications ChIP-seq data (Zhang *et al*, 2008a). Taken together, our data suggest that the effect exerted by each of the mH2A variants in blocking MET is probably mediated through distinct molecular mechanisms, all of which should be related to the constant mobility of mH2A1 variants from region to region, including promoters, to control gene expression.

### Composite Nanog/mH2A1.1 nucleosome elements define pluripotency

A surprising result derived from our experiments is that mH2A1.1-nucleosomes in ESCs are functionally linked to genes involved in mechanisms of pluripotency (Fig. EV1B, lower-right panel). To investigate the functional relevance of this observation, we focused on analyzing the average binding frequency of mH2A1.1-nucleosomes to all annotated transcription start sites (TSSs) across the genome during reprogramming. Figure EV4A shows that promoter-bound mH2A1.1-nucleosomes are dynamically redistributed by disappearing and reappearing at different elements during reprogramming. Interestingly, a significant number of the mH2A1.1-nucleosomes appearing in ESCs after the mobility phase (Days 1-9) bind to a novel set of promoter sites (see bracket in Fig. EV4A Group G_1.1_). The common promoter sites between MEFs and ESCs correspond to promoters associated with housekeeping genes (see Figs. EV4A bracket Group A_1.1_ and EV1B upper-left panel).

Next, we hypothesized that the novel Group G_1.1_ promoter-proximal sites emerging in ESCs might function as pluripotency-specific regulatory elements. To test this hypothesis, we assessed whether Group G_1.1_ elements are co-occupied by Nanog, the key pluripotency transcriptional regulator. Remarkably, we found that Nanog binding extensively overlaps with mH2A1.1 nucleosome locations in ESCs (Fig. 2A, left panel), whereas its binding overlap with mH2A1.2 sites is significantly lower (Fig. 2A, right panel). Reciprocally, Nanog binding was also found to be more robust, sharp and localized at mH2A1.1-bound elements as compared to mH2A1.2-bound sites (Fig. 2B). Pathway enrichment analysis of the genes bearing the composite mH2A1.1–Nanog bound sites revealed strong enrichment for pluripotency-related processes, including known pluripotency factors such as *Nanog*, *Pou5f1* (Oct4), *Sox2*, and *Myc* (Fig. 2C, left panel). In contrast, the commonly mH2A1.2–Nanog bound target genes displayed significantly lower enrichment for pluripotency genes (Fig. 2C, right panel).

**Figure 2:**
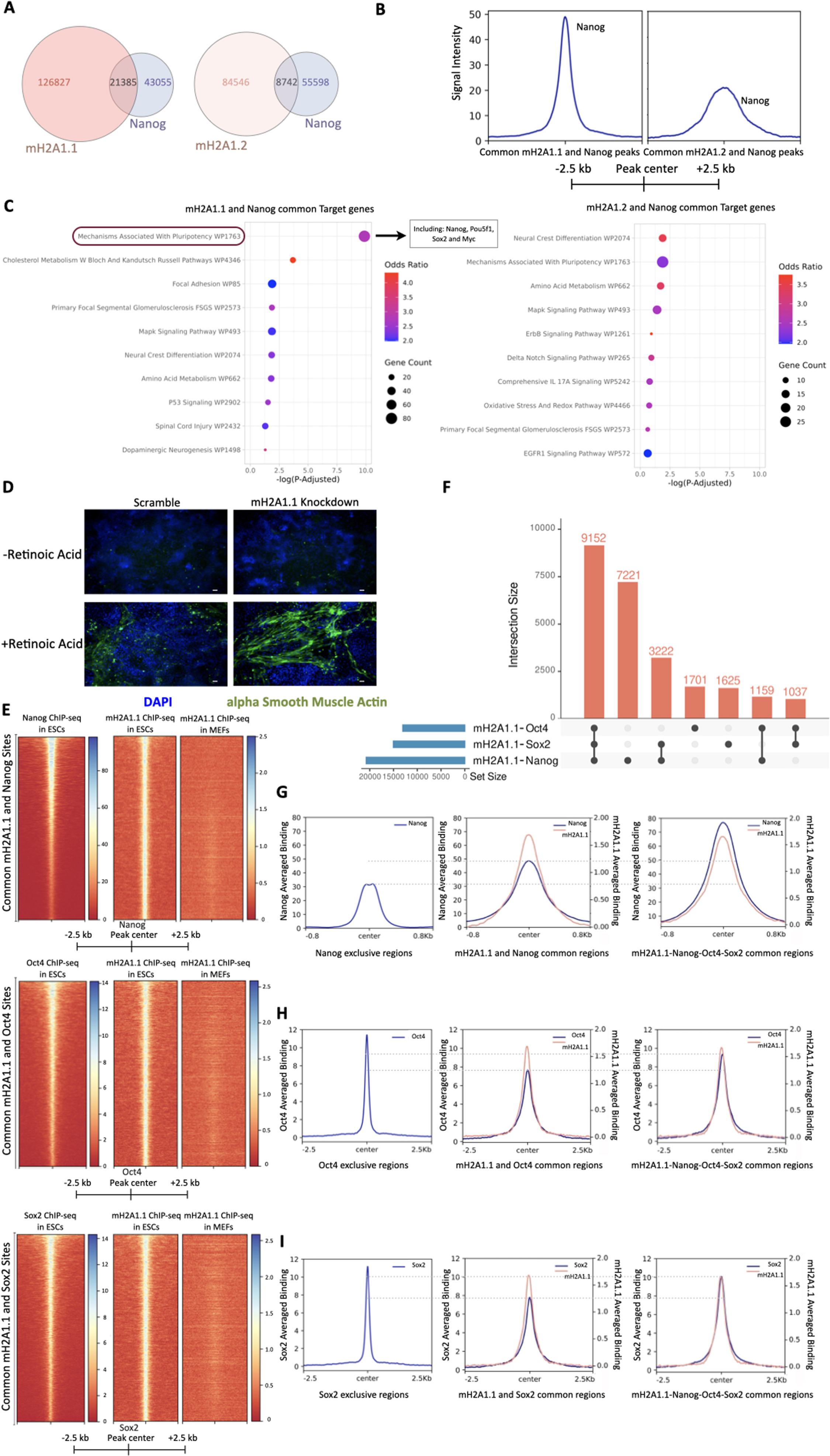
Composite Nanog/mH2A1.1 nucleosome elements protect pluripotency. (A) Venn diagrams showing the overlap of genomic regions co-occupied by Nanog and either mH2A1.1 (left) or mH2A1.2 (right) in ESCs. (B) Summary plots showing Nanog ChIP-seq signal (IP minus Input) across regions commonly co-bound by Nanog with either mH2A1.1 (left) or mH2A1.2 (right), centered on Nanog peaks and spanning ±2.5 kb. (C) Functional enrichment analysis (over-representation analysis, ORA) of genes near the Nanog/mH2A1.1 (left) and Nanog/mH2A1.2 (right) co-bound regions. Top 10 terms from the Wikipathways library are shown, ranked by adjusted p-value (P-Adjusted < 0.05). (D) Confocal images of ESCs treated with retinoic acid (lower panel) or untreated (upper panel), following lentiviral knockdown of mH2A1.1 (shRNA) or scramble control. Cells were fixed and stained with α-smooth muscle actin (α-SMA in green) to assess differentiation; nuclei counterstained with DAPI (in blue). Images acquired at 6days post-Retinoic Acid treatment. Scale bar: 40 μm. (E) Heatmaps depicting binding of Nanog (upper panel), Oct4 (middle panel), and Sox2 (bottom panel) to mH2A1.1 nucleosomes in ESCs, which are absent from MEFs, spanning ±2.5 kb from Nanog, Oct4 or Sox2 peak centers. (F) UpSet plot depicting overlapping regions between mH2A1.1 and Nanog, Oct4 or Sox2 and combinations thereof. The total number of mH2A1.1-nucleosome regions bound by any of the Oct4, Sox2 or Nanog is 25,117. The plot was created using Intervene(Khan & Mathelier, 2017). (G) Summary plots depicting the Averaged Signal (IP - Input) of Nanog (blue line) and mH2A1.1 (red line) in either Nanog exclusive regions (left panel), common mH2A1.1 and Nanog regions (middle panel) or common mH2A1.1-Nanog-Oct4-Sox2 regions (right panel). The horizontal gray dashed lines compare the peaks of Nanog signal among the three classes of regions. Common regions were defined by the UpSet plot (F) and were downsampled to acquire equal number of regions analyzed per region condition. (H) Same as (G) but for Oct4 binding (I) Same as (G) but for Sox2 binding

The above results suggest that mH2A1.1 nucleosomes play a functional role in pluripotency. To address this, we analyzed the transcriptome of ESCs following knockdown (KD) of either mH2A1.1 or mH2A1.2 using our shRNA lentivirus system (Valakos *et al*, 2023). In agreement with our chromatin-based observations, we found that mH2A1.1 KD—but not mH2A1.2 KD—led to a downregulation of key pluripotency TFs including *Nanog*, *Klf4* and *Esrrb* (Fig. EV4B) and genes related to “Mechanisms Associated With Pluripotency” (Fig. EV4C). Furthermore, 35% of the DEGs in mH2A1.1 KD ESCs are direct targets of Nanog (Fig. EV4D) associated with Pluripotency Pathways (Fig. EV4E). The biological significance of the results described above was demonstrated by showing that mH2A1.1 KD ESCs display enhanced differentiation capability upon retinoic acid treatment as compared to control cells, thus indicating that in the absence of mH2A1.1, the pluripotent phenotype is less stable (Fig. 2D).

By plotting ChIP-seq signals, we discovered that Nanog binds with high affinity to sites contained within mH2A1.1 nucleosomes (Fig. 2E, upper panel). Strikingly, these composite Nanog-mH2A1.1-bound chromatin elements are novel mH2A1.1 nucleosome sites since they are not occupied by mH2A1.1 in MEFs, despite its abundance in these cells (Fig. 2E, upper panel). Accordingly, the novel Nanog-mH2A1.1 nucleosome regions are highly enriched for active chromatin marks, such as H3K27ac, H3K9ac, H3K4me1 and H3K4me2 in ESCs, but not in MEFs (Figs. EV4F, G, H, I, respectively), further supporting their functional relevance to gene expression. Given that Nanog forms Transcription Factor hubs with Oct4 and Sox2 in maintaining pluripotency (Kim *et al*, 2008; Orkin & Hochedlinger, 2011), we asked whether the ESC-specific mH2A1.1-bound regulatory elements are also co-occupied by Oct4 and Sox2. We found that in contrast to the preference of Nanog binding to mH2A1.1 nucleosomes (Fig. 2G, compare left with middle panel), Oct4 or Sox2, are not preferentially associated with mH2A1.1 nucleosomes in ESCs (Figs. 2H and 2I, left and middle panels), although they do co-bind with mH2A1.1 specifically in ESCs (Fig. 2E, middle and lower panels). Strikingly, however, all three pluripotency-specific TFs assemble hubs on mH2A1.1 nucleosomes in ESCs, with Nanog binding being further increased at these composite elements (Fig. 2F and Figs. 2G, H, I right panels).

These data strongly support a model in which mH2A1.1 plays a central role in building a landscape suitable for the assembly of pluripotency TF hubs, thus functioning as epigenetic bookmark to establish and/or stabilize stemness networks through its selective recruitment to promoters of pluripotency-associated genes (see model in Fig. 6C).

### Temporal dynamics and repositioning of mH2A1.2 nucleosomes during reprogramming

The above findings suggest that the massive genome-wide rearrangement of the mH2A1-nucleosome landscape during reprogramming, is functionally linked to the activation of transcriptional regulatory networks necessary for pluripotency. To further investigate this unexpected finding, we focused on the mH2A1.2-enriched Group C_1.2_ regions, because they are characterized by sharply defined nucleosome positioning patterns, suggesting nucleosome phasing at the same genomic positions across different time-points (Fig. 1E). We first tested Group C_1.2_ regions for OSKM binding and found a strong correlation between OSKM binding and Group C_1.2_ occupancy (Fig. 3A). This observation aligns with the finding that Group C_1.2_ regions (1190 regions) exhibit the highest promoter enrichment and chromatin accessibility among all mH2A1-bound genomic elements (Figs. 1F-G). Although a general correlation between OSKM-bound sites and mH2A1.2 nucleosome localization is lacking (Fig. 1A right panel), Group C_1.2_ regions represent a striking notable exception (Fig. 3A).

**Figure 3:**
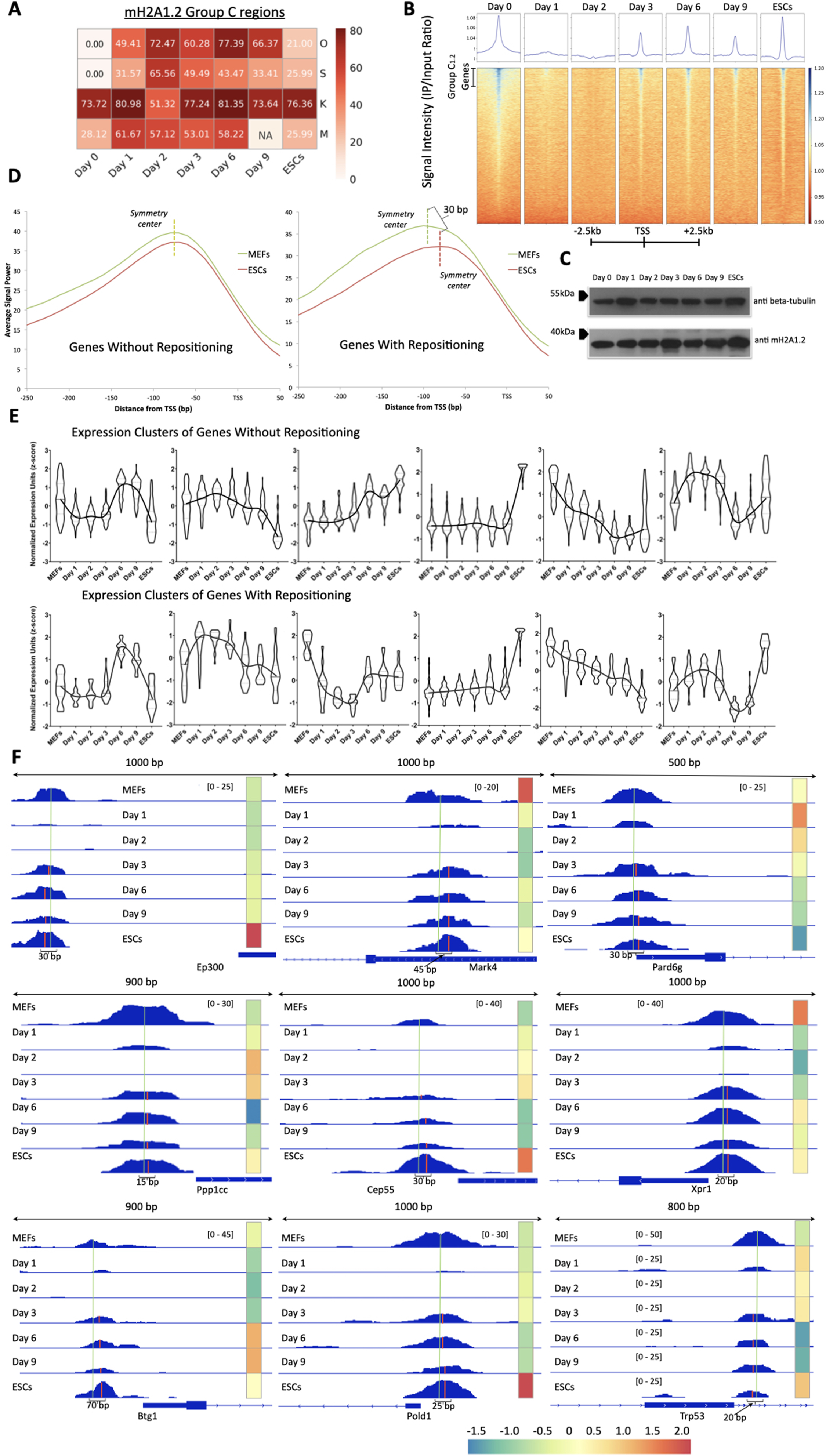
Temporal dynamics and repositioning of mH2A1.2 nucleosomes during reprogramming. (A) Heatmap showing the percentage of shared binding peaks between each of the O/S/K/M factors (indicated on the right) and the mH2A1.2-bound Group C_1.2_ regions (as identified in Fig. 1E) across Day 0, reprogramming stages (Day 1, Day 2, Day 3, Day 6, Day 9) and ESCs. NA: Not Available (B) Summary plots and heatmaps displaying mH2A1.2 ChIP-seq signal (IP/Input ratio) across all annotated mouse promoters from –2.5 kb to +2.5 kb relative to the TSS of all genes at Day 0, throughout reprogramming (Day 1, Day 3, Day 6, Day 9) and ESCs. Regions were sorted in descending order based on IP signal at MEFs (Day 0) state, with the same order maintained across time-points. Signal intensities are shown as IP/Input ratio. Notably, mH2A1.2 nucleosomes transiently disappear from transcription start sites (TSSs) on Day 1 and reappear by Day 3 at the same loci. (C) Western blot analysis showing mH2A1.2 protein levels across reprogramming time-points and in ESCs. β-tubulin was used as a loading control. mH2A1.2 protein expression remains relatively stable throughout the process. (D) Summary plots illustrating comparison of the nucleosome occupancy between genes with stable mH2A1.2 peak positions (left panel) and genes with nucleosome repositioning (right panel). mH2A1.2 nucleosome repositioning is characterized by an average 30 bp shift in peak center between MEFs and ESCs (compare green and red dashed lines in right panel). (E) Violin plots comparing expression dynamics (z-score) of gene clusters with either stable (top) or repositioned (bottom) mH2A1.2 nucleosomes throughout reprogramming. Median expression trends across time-points are connected by lines. RNA-seq data were derived from (Valakos *et al*, 2023) and (Klagkou *et al*, 2024). (F) IGV browser snapshots of mH2A1.2 ChIP-seq bigwig tracks in MEFs (Day 0), during reprogramming (Days 1–9), and in ESCs for representative genes undergoing nucleosome repositioning. All tracks were normalized by subtracting Input from the IP signal. A color bar indicates z-score normalized gene expression across time-points. The green line marks the nucleosome symmetry center at Day 0; red lines mark new symmetry centers in repositioned states at subsequent time-points.

We then calculated the average binding frequency of mH2A1.2 nucleosomes at all annotated TSSs (Fig. 3B). We discovered that mH2A1.2 nucleosomes corresponding to ∼1,000 gene promoters and proximal regulatory elements of Group C_1.2_ (Fig. 1E) are evicted as early as Day 1 of reprogramming and re-deposited by Day 3 at the same locations, where they remain stably bound throughout the remainder of the reprogramming timeline (Fig. 3B). Importantly, this disappearance and reappearance is not due to degradation and resynthesis of the mH2A1.2-protein, as its overall levels remain relatively stable throughout reprogramming (Fig. 3C).

Since well-positioned, promoter-bound nucleosomes—such as those found in Group C_1.2_ regions—are known to regulate TF binding (Agelopoulos & Thanos, 2006; Lavigne *et al*, 2015; Nocetti & Whitehouse, 2016; Rudnizky *et al*, 2019), we investigated whether the disappearance and reappearance of mH2A1.2 promoter nucleosomes during reprogramming has functional consequences. To this end, we mapped the precise positions of Group C_1.2_ nucleosomes in MEFs (prior to their disassembly) and in ESCs (following their reassembly), representing the start and end points of reprogramming. Nucleosome occupancy was defined as the probability of a base pair being occupied by a nucleosome. Strikingly, our analysis revealed a subset of 255 genes in which mH2A1.2 nucleosomes reappear at positions slightly different from their original location in MEFs (Dataset EV4). The average positional shift of this specific group of nucleosomes was calculated to be ∼30bp (Fig. 3D, right panel). In contrast, the majority of Group C_1.2_ genes (792 genes) showed reappearance of mH2A1.2-nucleosomes at the exact same nucleotide position as before their disassembly (Fig. 3D left panel, Dataset EV4).

To assess the functional implications of nucleosome repositioning, we compared gene expression patterns between these two groups. Surprisingly, we did not observe a global difference between their gene expression profiles during reprogramming (Fig. 3E), a finding consistent with the OSKM DNA binding profile (Fig. EV2E). However, further gene-by-gene analysis throughout the reprogramming timeline revealed many individual cases where nucleosome repositioning correlated with distinct expression dynamic changes of the underlying genes (Fig. 3F). For instance, p300, a histone acetyltransferase and global transcriptional co-factor, showed a progressive increase in expression after the reappearance of the mH2A1.2 nucleosome to its new position 30 bp upstream from the original site, reaching maximal expression in ESCs. Similar trends were observed for the *Pold1* and *Btg1* genes, both of which exhibited increased expression correlated with nucleosome repositioning. In contrast, *Mark4* and *Pard6g* displayed a progressive decline in their expression, corresponding to the repositioned nucleosomes towards transition to pluripotency. Moreover, genes such as *Ppp1cc*, *Trp53* and *Cep55* exhibited fluctuating expression levels that coincided with dynamic repositioning of mH2A1.2 nucleosomes over time.

Taken together, these findings identify a novel, dynamically reprogrammable fraction of chromatin, wherein mH2A1.2 nucleosomes are re-deposited at new positions, correlating with context-specific alterations in gene expression. This positional plasticity of histone variant nucleosomes may represent an additional layer of epigenetic regulation during cell fate transitions.

### Repositioning of mH2A1.2 nucleosomes modulates E2F4 binding and gene expression during reprogramming

Previous studies have shown that well-positioned nucleosomes at promoter regions can modulate the accessibility of TFs and the general transcriptional machinery to their target DNA sequences (Lomvardas & Thanos, 2001; Lavigne *et al*, 2015; Nocetti & Whitehouse, 2016; Rudnizky *et al*, 2019). Given our observation that mH2A1.2-nucleosome relocation often correlates with altered gene expression, we hypothesized that these mH2A1.1 nucleosome positional changes might also impact TF binding dynamics during reprogramming. To investigate this, we first performed GO analysis on the subset of genes exhibiting mH2A1.2-nucleosome repositioning and found that the majority are involved in cell-cycle regulation and cell division—both key processes for successful reprogramming (Fig. 4A). Notably, most of these genes were lowly expressed or repressed in MEFs but became upregulated in ESCs (Fig. 4B) supporting the idea that mH2A1.2-nucleosome positioning might regulate TF access to these loci leading to activation of their expression in ESCs.

**Figure 4:**
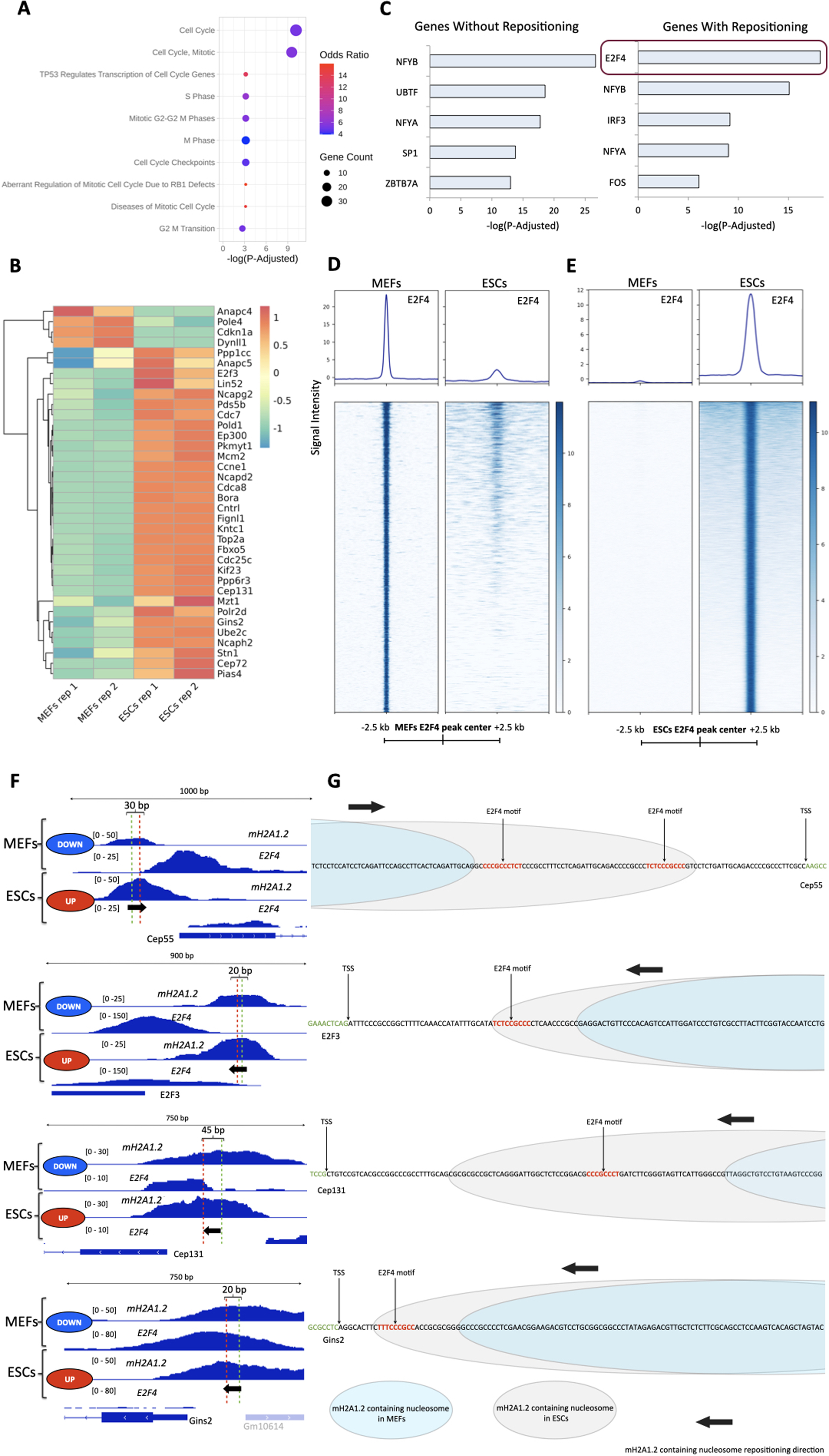
mH2A1.2 nucleosome repositioning switches on or off E2F4 DNA binding during reprogramming. (A) Pathway enrichment analysis (Reactome) of genes exhibiting mH2A1.2 nucleosome repositioning between MEFs and ESCs. Cell cycle- and division-related pathways are significantly enriched. Shown are the top 10 enriched terms based on adjusted p-values (P-Adjusted < 0.05). (B) Heatmap displaying expression levels (z-score) of cell cycle-related genes (from Fig. 4A) that undergo mH2A1.2 nucleosome repositioning in ESCs compared to MEFs. RNA-seq data were derived from (Valakos *et al*, 2023) and (Klagkou *et al*, 2024). (C) Transcription factor target enrichment analysis (TRUSST Database) comparing genes without mH2A1.2 repositioning (left) to those with repositioning (right). Genes with repositioned nucleosomes show specific enrichment for E2F4 target genes. Top 5 terms are shown, sorted by P-Adjusted value < 0.05. (D) Summary and heatmap plots depicting strong E2F4 binding in MEFs (left panel) and reduced E2F4 binding to the same genomic regions in ESCs. Signal Intensity was computed by subtracting Input from the IP signal. (E) Same as (D), but comparing E2F4 binding sites in ESCs (right) to the same regions in MEFs (left). (F) IGV browser snapshots of ChIP-seq tracks showing mH2A1.2 and E2F4 binding at representative genes undergoing mH2A1.2 nucleosome repositioning. All tracks were normalized by subtracting Input from the IP signal. Black arrows indicate the direction of nucleosome repositioning, with green and red dashed lines marking original and final nucleosome symmetry centers, respectively. Repositioning inhibits E2F4 DNA binding in ESCs, facilitating transcriptional activation of cell cycle-promoting genes. Side ovals indicate relative gene expression (blue: low, red: high). (G) Schematic representation illustrating the repositioning mechanism: in MEFs, mH2A1.2 nucleosomes (light blue ovals) do not obstruct E2F4 motifs (red text in DNA sequence), allowing E2F4 binding and transcriptional repression. In ESCs, repositioned mH2A1.2 nucleosomes (gray ovals) occlude E2F4 motifs, blocking E2F4 access and enabling gene activation.

Therefore, we conducted a comparative Transcription Factor Target Enrichment analysis between genes with and without mH2A1.2 repositioning and found that genes with repositioned nucleosomes were significantly enriched for E2F4 targets (Fig. 4C, right panel). The TF E2F4 is a well-known negative regulator of cell-cycle progression and a key player in both tumor suppression and pluripotency regulation (Trimarchi *et al*, 2001; Dimova & Dyson, 2005; Chen *et al*, 2009; Kent & Leone, 2019; Hsu *et al*, 2019). Consistent with this, the E2F4 DNA-binding motif was the fourth top enriched motif in the regulatory regions of genes undergoing nucleosome repositioning, but it was absent from genes without mH2A1.2-nucleosome repositioning (Figure EV5A). Therefore, we hypothesized that the differential positioning of mH2A1.2-nucleosomes before (MEFs) and after completion of reprogramming (ESCs) could modulate E2F4 access to its binding sites.

To test this, we performed ChIP-seq for E2F4 in both MEFs and ESCs and found that E2F4 binds distinct genomic loci in MEFs and ESCs, with minimal overlap (Figs. 4D and 4E). This suggests that despite its ubiquitous expression, E2F4 target sites are cell type-specific, thus implying a regulatory mechanism that could be governed by chromatin context. Functional enrichment analysis of E2F4 target genes in MEFs and ESCs confirmed their enrichment in cell-cycle–related pathways (Figs. EV5B, C). To directly assess the impact of mH2A1.2-nucleosome positioning on E2F4-binding motifs, we integrated our topological maps of mH2A1.2-nucleosome occupancies with E2F4-binding sites. Figure 4F highlights several examples demonstrating that repositioning of mH2A1.2-nucleosomes correlates with a gain or loss of E2F4 binding at target promoters. In ESCs, E2F4-occupancy is significantly reduced at these sites due to the masking of E2F4-binding motifs by mH2A1.2-nucleosome repositioning leading to derepression and transcriptional activation (Figs. 4F, G). For example, at the *Cep55* promoter, a 30 bp mH2A1.2-nucleosome positioning shift results in occlusion of two E2F4 binding motifs, preventing binding and thereby lifting E2F4-mediated repression (Figs. 4F, 4G and EV5F). Similar patterns were observed at the promoters of *E2F3* (positive regulator of the cell cycle, 20 bp shift), *Cep131* (involved in centrosome duplication, 45 bp shift), *Gins2* (required for initiation of DNA replication, 20 bp shift), *Cdc7* (a cell cycle division protein, 50 bp shift) and *Pds5b* (regulatory subunit of the cohesin regulator, 30 bp) (Figs. 4F, G and EV5D, E, F). In all cases, the repositioned mH2A1.2-nucleosomes prevent E2F4 binding, thus allowing activation of genes critical for cell-cycle progression and thereby supporting the transition towards pluripotency. Importantly, we validated these findings by demonstrating that the mH2A1.2-nucleosome-repositioning effect at various target genes is similar between ESCs and iPSCs (Fig. EV5F).

Taken together, these findings demonstrate a cell type–specific interplay between mH2A1.2-nucleosome positioning and E2F4 binding at promoters of genes involved in cellular reprogramming (Fig. 6B). This mechanism underscores the functional relevance of nucleosome repositioning as an epigenetic switch, dynamically regulating TF access to shape cell fate decisions.

### NRF-1 and mH2A1.2 coexist in repositioned nucleosomes in ESCs

To address the mechanisms driving mH2A1.2-nucleosome repositioning at specific promoters during reprogramming, we analyzed the DNA sequences associated with mH2A1.2-occupancy. We have previously shown that the bZIP transcription factor NRF-1 frequently co-occupies the same DNA regions as mH2A1.2 nucleosomes, forming highly stable composite NRF-1/mH2A1.2-nucleosome complexes that are enriched at promoters of highly expressed genes, thereby enhancing transcriptional robustness (Lavigne *et al*, 2015). Interestingly, we found that NRF-1 expression is upregulated in ESCs (Fig. 5A), suggesting a potential cell-type–specific role for NRF-1 in guiding mH2A1.2 nucleosome relocation.

**Figure 5:**
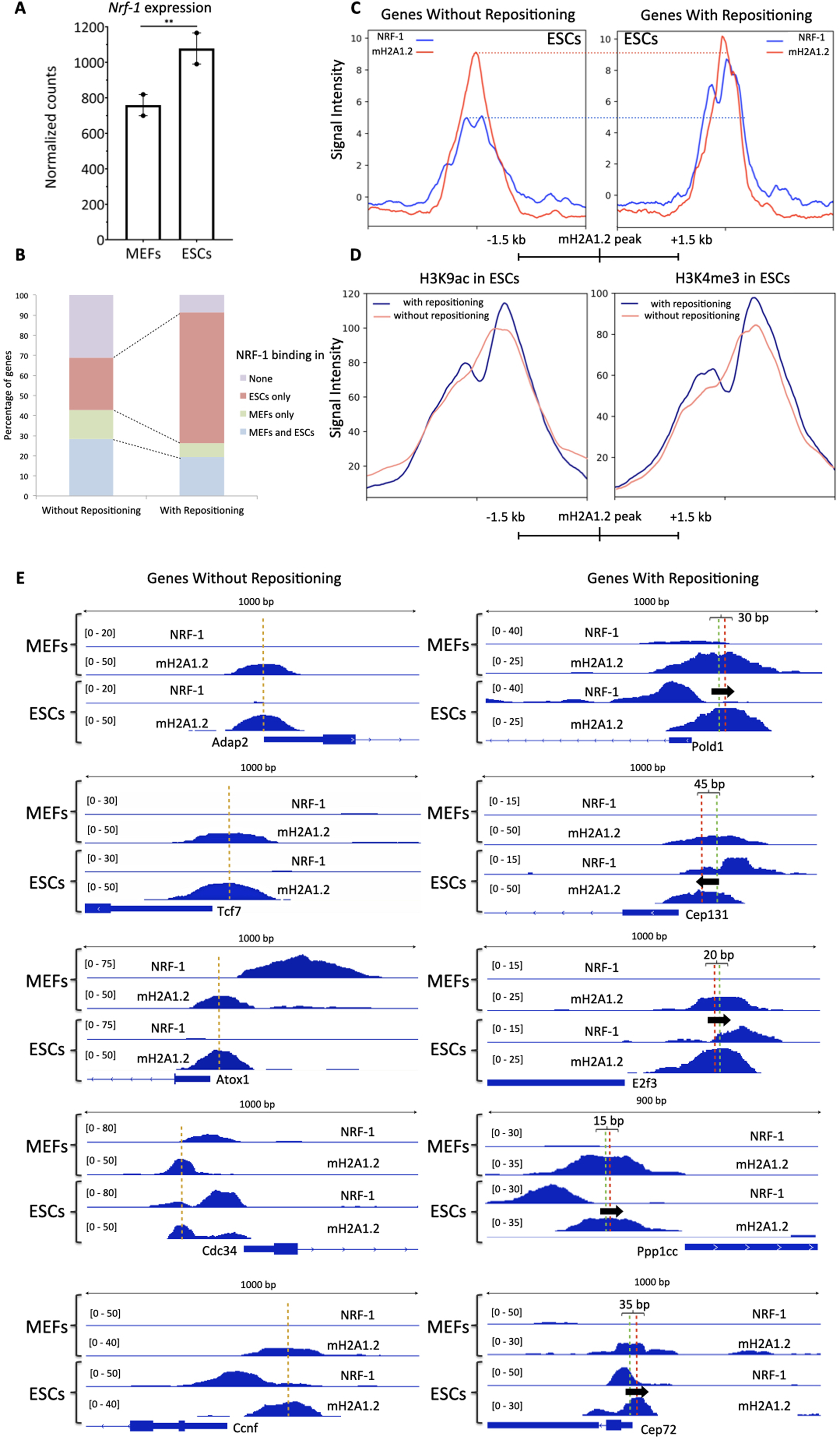
Specific NRF-1 binding in ESCs coincides with mH2A1.2 nucleosome repositioning. (A) Bar graph depicting normalized RNA expression levels of *Nrf-1* in MEFs and ESCs. Data are presented as mean ± SEM (N = 2). RNA-seq datasets were retrieved from (MEFs)^49^ and (ESCs)(Valakos *et al*, 2023). Statistical significance was defined with DESeq2 (Klagkou *et al*, 2024). **: P-Adjusted<0.01 (B) Bar graph showing the percentage of genes without (left) and with (right) mH2A1.2 nucleosome repositioning bound by NRF-1 as indicated. (C) Summary plots depicting mH2A1.2 and NRF-1 binding at genomic regions neighboring to genes without (left) and with (right) mH2A1.2 nucleosome repositioning as indicated. ChIP-seq signal was normalized by subtracting Input from IP. X Axis corresponds to the distance from mH2A1.2 peak center. ChIP-seq data for NRF-1 were retrieved from (Baird *et al*, 2017) and (Domcke *et al*, 2015). The horizontal dashed lines compare the ChIP-seq signal for both mH2A1.1 and NRF-1 between the two classes of genes. (D) Summary plots depicting H3K9ac (left panel) and H3K4me3 (right panel) signal in ESCs, for genes with nucleosome repositioning (dark blue line) and without repositioning (pale red line). X Axis corresponds to the distance from mH2A1.2 peak center. ChIP-seq signal was normalized by subtracting Input from IP. Data for H3K9ac and H3K4me3 were retrieved from (Chronis *et al*, 2017). (E) IGV browser snapshots of representative genes depicting distinct NRF-1 binding patterns between genes without (left) and with (right) mH2A1.2 nucleosome repositioning. For genes without repositioning, examples for each NRF-1 binding category from (B) is shown (from top left to bottom left: no binding (2 examples), MEF-only binding, binding to both MEFs and ESCs, ESCs-only binding. Yellow dashed lines indicate nucleosome symmetry center in both MEFs and ESCs. For genes with the mH2A1.2 repositioned nucleosome, black arrows show the direction of nucleosome movement; green and red dashed lines mark the original and final peak centers. NRF-1 binding in ESCs is associated with repositioned mH2A1.2 nucleosomes. Nucleosome repositioning distances were calculated from peak center shifts in mH2A1.2 ChIP-seq data.

**Figure 6:**
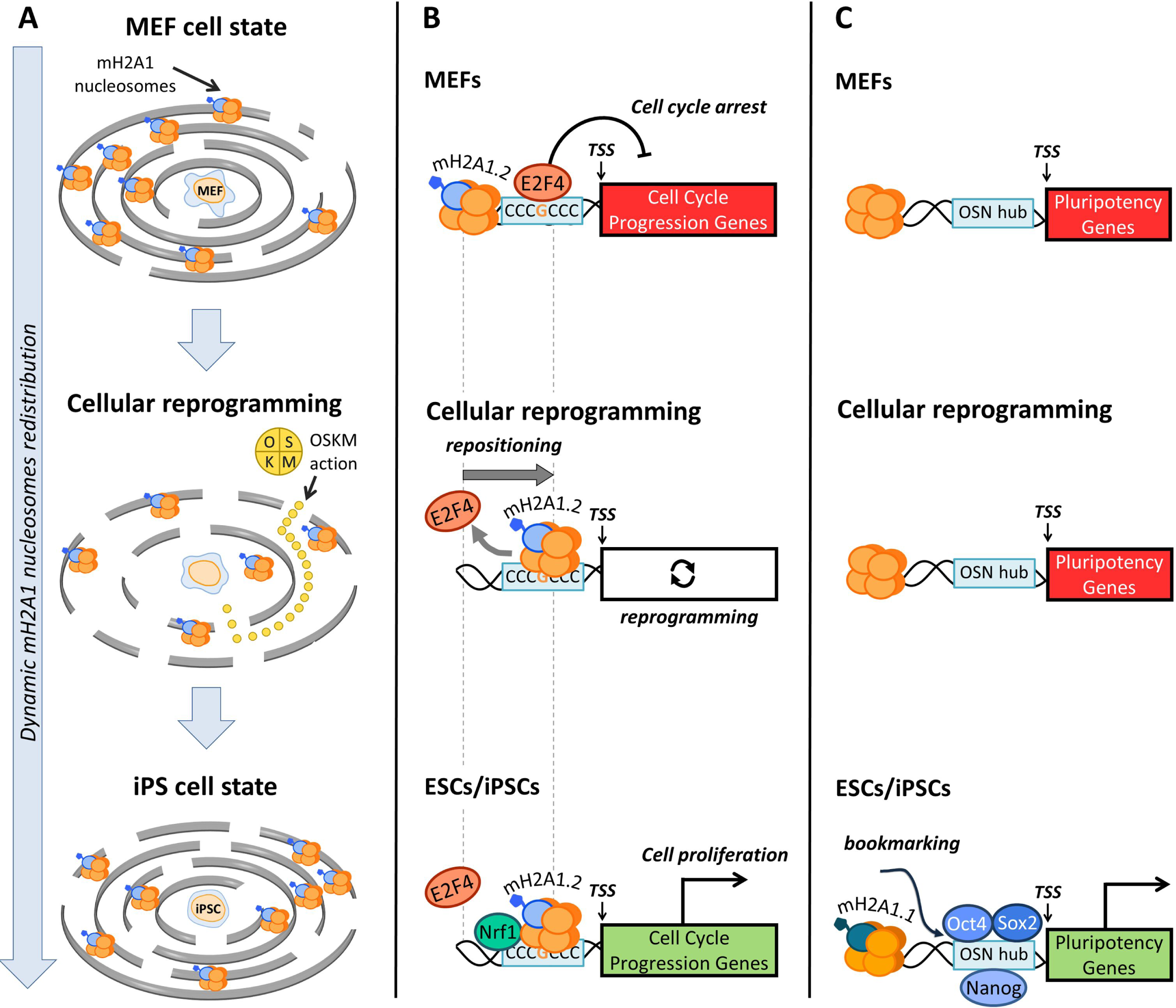
Models depicting the pleiotropic role of mH2A1 nucleosomes in cellular reprogramming. (A) mH2A nucleosomes function as epigenetic barriers safeguarding the cell identity of somatic cells by stabilizing cell type-specific gene expression programs. (B) Proposed model depicting that in MEFs, E2F4 binds to and represses cell cycle-promoting genes, inducing cell-cycle arrest. In ESCs/iPSCs, mH2A1.2 nucleosome repositioning occurred during reprogramming and facilitated by the specific NRF-1 binding—blocks E2F4 access, leading to transcriptional activation of cell cycle-promoting genes and induction of cell proliferation. (C) Model depicting that in ESCs, mH2A1.1-containing nucleosomes occupy promoter regions that are direct targets of Nanog, Oct4 and Sox2. These regions are devoid of mH2A1.1 in MEFs, underscoring the role of mH2A1.1 nucleosome reconfiguration during reprogramming and highlight its role as a position-dependent epigenetic gatekeeper of pluripotency.

To test this, we analyzed NRF-1 ChIP-seq data comparing NRF-1-occupancy between promoters of genes with and without repositioned mH2A1.2 nucleosomes in MEFs and ESCs. Strikingly, we observed a strong and preferential binding of NRF-1 to genes exhibiting mH2A1.2 nucleosome repositioning (65% of the repositioned genes as opposed to 26% in non-repositioned), specifically in ESCs as compared to MEFs (Fig. 5B, Dataset EV4). This preferential occupancy aligns with the elevated NRF-1 expression in ESCs suggesting that NRF-1 may facilitate or stabilize the repositioning of mH2A1.2-nucleosomes. Moreover, NRF-1-binding was generally stronger and more localized at repositioned promoters in ESCs, consistent with the generation of a precisely organized and stable chromatin environment (Fig. 5C). Supporting this, we detected slightly elevated levels of H3K9ac and H3K4me3 with sharper signals in these regions, further indicative of a transcriptionally active and robust chromatin configuration of the repositioned nucleosomes in ESCs (Fig. 5D).

Figure 5E depicts representative examples illustrating the precise architecture of the composite NRF-1/mH2A1.2-nucleosome complex in both repositioned and non-repositioned genes in MEFs and ESCs. Remarkably, in the vast majority of the genes undergoing mH2A1.2-nucleosome repositioning, NRF-1-binding was absent in MEFs with the mH2A1.2-nucleosomes occupying their original positions, while in ESCs, NRF-1 binds robustly and the mH2A1.2-nucleosome is consistently repositioned. In contrast, in non-repositioned genes, the mH2A1.2-nucleosome maintains the same stable position across both cell types, and NRF-1-binding is variable—either absent in both cell types, present only in MEFs or ESCs, or present in both (Fig. 5E).

Taken together, these findings suggest a functional interplay between NRF-1 binding and mH2A1.2-nucleosome repositioning, wherein NRF-1 may facilitate or stabilize the new position of the mH2A1.2 nucleosomes during reprogramming. This cell type–specific cooperative interaction likely contributes to the establishment of transcriptionally robust chromatin states at critical pluripotency regulatory genes in ESCs.

## Discussion

We show that the histone variant mH2A1 exerts a stage-specific functional repurposing during cellular reprogramming. At the beginning of reprogramming, mH2A1 primarily acts as a barrier to cellular plasticity by actively inhibiting MET, an essential early event for initiating the conversion of somatic to pluripotent cells (Pasque *et al*, 2011; Gaspar-Maia *et al*, 2013; Barrero *et al*, 2013; Pliatska *et al*, 2018; Valakos *et al*, 2023). In addition to these earlier observations that assigned narrowly defined roles to mH2A1 nucleosomes, our findings indicate that this histone variant is repurposed into a facilitator of reprogramming via its dynamic repositioning on chromatin contributing to both establishing and maintaining the pluripotent state (Fig. 6A).

This dual-function aligns with a broad evolutionary principle involving the functional repurposing of many molecular components allowing individual structural or regulatory components to acquire distinct roles in different contexts rather than being fixed in one mode of action. Previous studies have shown that in the epigenetic and chromatin context, the Polycomb repressive complexes (PRC1/PRC2), classically viewed as transcriptional silencers, are repurposed during certain developmental stages and in pluripotent cells to facilitate enhancer activation and chromatin looping, functions that depend on altered complex composition (Cruz-Molina *et al*, 2017; Hong *et al*, 2024). Likewise, the histone variant H3.3 in differentiated cells marks heterochromatin and stabilizes repressed domains, but in pluripotent cells and active chromatin regions, it marks transcribed genes and enhancers (Schlesinger *et al*, 2017; Tafessu *et al*, 2023). Notably, the pluripotency transcription factor NANOG is repurposed from an activator to a transcriptional repressor to repress the expression of SOX2 before implantation (Wong *et al*, 2025). These context-dependent functional reversals, driven by changes in nucleosome dynamics and deposition machinery, mirror the dual and repurposed function of mH2A1 during reprogramming.

The pleiotropic functions of mH2A1 arise from its dynamic genome-wide reconfiguration during reprogramming, which enables precise control of transcription factor (TF) access to chromatin. Although mH2A1 variants were originally viewed as roadblocks to pluripotency (Pasque *et al*, 2011; Gaspar-Maia *et al*, 2013; Barrero *et al*, 2013), our findings show that this interpretation reflects only their early role in maintaining somatic cell identity. At the onset of reprogramming, extensive eviction and repositioning of mH2A1 nucleosomes help cells overcome constraints such as E2F4-mediated cell-cycle arrest and the mH2A1-dependent mesenchymal gene-expression network. This large-scale nucleosome mobility also underlies the emergence of heterogeneous intermediate states typical of early iPSC formation (Apostolou & Stadtfeld, 2018; Schiebinger *et al*, 2019), reflecting transient release of pluripotency-related genes from repression.

Our data identify mH2A1.2 as a central driver of functional repurposing during reprogramming (Fig. 6B). In somatic MEFs, mH2A1.2 nucleosomes enforce the mesenchymal phenotype by restricting E2F4 binding on cell-cycle gene promoters (Ni *et al*, 2020). Early in reprogramming, ∼1,000 genes with phased mH2A1.2 nucleosomes lose these nucleosomes and by Day 3, 255 regain them at positions shifted by ∼30 bp. In their new locations, these nucleosomes occlude E2F4 binding, reversing their original function and promoting the robust cell-cycle activity required for transition toward pluripotency. This nucleosome repositioning is guided, at least in part, by NRF-1, which co-localizes with and may anchor the newly positioned nucleosomes through interactions with mH2A1.2 (Gamarra & Narlikar, 2021; Sun & Bernstein, 2019). These findings reveal a reversible, context-dependent nucleosome-positioning switch whose transcriptional outcome depends on timing, chromatin environment, and cooperating TFs.

In contrast, mH2A1.1 functions as a chromatin bookmark that supports pluripotency maintenance (Fig. 6C). In ESCs, mH2A1.1-containing nucleosomes are assembled at newly established regulatory sites enriched for pluripotency genes and promote the formation of Nanog, Oct4, and Sox2 TF hubs (Kim *et al*, 2008). Loss of mH2A1.1 increases the ESCs’ differentiation potential, consistent with its role in stabilizing pluripotency-related transcriptional programs. Its bookmarking activity may stem from its ability to bind ADP- and poly-ADP-ribose. We propose that the stable assembly of mH2A1.1 at these sites represents a repurposed late-stage function building a flexible but stable pluripotent regulatory architecture.

Together, these findings show how mH2A1 isoforms fine-tune nucleosome positioning so the genome is neither too rigid to block state transitions nor too permissive to risk instability. Through sequential functional repurposing—first enforcing somatic identity, then enabling cell-cycle entry, and finally shaping pluripotency architecture—, mH2A1 switches from a reprogramming brake to a coordinator of the steps leading to pluripotency. This dynamic adaptability exemplifies a broader evolutionary principle: the reuse of a single chromatin component to perform distinct, context-dependent functions. mH2A1’s versatility highlights how chromatin regulators act not as fixed determinants of cellular fate but as flexible agents of plasticity, tuned to the needs of the cell’s developmental trajectory.

## Methods

### Cell lines and mouse embryonic fibroblasts isolation

Bruce 4 (C57BL/6 strain) mouse Embryonic Stem Cells (mESCs) and Human Embryonic Kidney 293T (HEK293T) were purchased from ATCC. Mouse Embryonic Fibroblasts (MEFs) were isolated by E13.5 C57BL/6 mouse embryos, which were surgically removed from pregnant female mice and placed in PBS (1X). Briefly, the embryos were dissociated, the head and the embryonic internal organs were removed from the abdominal cavity and then trypsinized and passed through a 10-ml syringe to produce single-cell suspensions which were expanded. MEFs and HEK293T were cultured in Dulbecco’s Modified Eagle’s Medium (Sigma-Aldrich, cat. D6429) supplemented with 10% heat-inactivated fetal bovine serum and 1x GlutaMAX™ Supplement (Gibco™, 35050-061), 1x MEM Non-Essential Amino Acids Solution (Gibco™, 11140-050) and Penicillin-Streptomycin mix diluted at 1/100 (Gibco™, 15140-122) at 37°C with 5% CO_2_. Bruce 4 mESCs were cultured in Knock Out Dulbecco’s Modified Eagle’s Medium (Gibco, 10829018), supplemented with 20% ES tested fetal bovine serum (Pansera, P30-2602), 1x GlutaMAX™ Supplement (Gibco™, 35050-061), 1x MEM Non-Essential Amino Acids Solution (Gibco™, 11140-050), Penicillin-Streptomycin mix diluted at 1/100 (Gibco™, 15140-122), 0.1 mM beta-mercaptoethanol (Sigma Aldrich, M6250) and with 20 ng/mL mLIF (Santa Cruz Biotechnology, sc-4378) at 37°C with 5% CO_2_.

All animals were handled according to the ethical standards of BRFAA under protocols approved by the Animal Care and Use Committee of the Institution and in accordance with the European Convention 123/Council of Europe and Directive 86/609/EEC. Animal experimental protocol was approved by an authorized veterinarian committee in accordance to Greek legislation (Presidential Decree 56/2013, in compliance with the European Directive 2010/63). Serial number of approval: 172227/08/02/2024.

### MEFs cellular reprogramming

For the cellular reprogramming experiments, we used mouse embryonic fibroblasts (MEFs) from cross-bred homozygous transgenic mice originating from the B6;129S4-Col1a1^tm1(tetO-Pou5f1,-Klf4,-Sox2,-Myc)Hoch^/J (RRID:IMSR_JAX:011001) and B6.Cg-Gt(ROSA)26Sor^tm1(rtTA*M2)Jae^/J (RRID:IMSR_JAX:006965) strains (Hochedlinger *et al*, 2005; Stadtfeld *et al*, 2010). MEFs were seeded in 10 cm plates and doxycycline was added (2 μg/ml final concentration) for the pluripotency factors induction until day 25. After Day 6, the cultures were maintained in iPSC medium [High glucose DMEM, 20% KnockOut Serum Replacement (10828028, Thermo Fisher Scientific), 1x GlutaMAX™ Supplement (Gibco™, 35050-061), 1x MEM Non-Essential Amino Acids Solution (Gibco™, 11140-050) and Penicillin-Streptomycin mix diluted at 1/100 (Gibco™, 15140-122), 0.1 mM beta-mercaptoethanol (Sigma Aldrich, M6250) with 20 ng/mL mLIF (Santa Cruz Biotechnology, sc-4378) at 37°C with 5% CO_2_.

### MNase-sequencing

Isolation of mono-di-tri nucleosomes and immunoprecipitation were performed as previously described (Lavigne *et al*, 2015). Briefly, cells were resuspended in 8ml of N-ChIP buffer I (0.3M Sucrose, 60mM KCl, 15mM NaCl, 5mM MgCl_2_, 0.1mM EGTA, 15mM Tris-HCl, pH 7.5, 0.5mM DTT). Next, lysis was performed by adding 2 mL of N-ChIP buffer II (0.3M sucrose, 60 mM KCl, 15mM NaCl, 5mM MgCl_2_, 0.1mM EGTA, 15mM Tris-HCl pH 7.5, 0.4% NP-40 ,0.5mM DTT) and the nuclei were purified with a sucrose cushion by layering lysed cells on 8 ml of N-ChIP-buffer III (1.2M sucrose, 60 mM KCl, 15mM NaCl, 5mM MgCl_2_, 0.1mM EGTA ,15 mM Tris-HCl, pH 7.5, 0.5mM DTT) followed by centrifugation at 3,000 rpm for 30 minutes. The supernatant was discarded and the nuclear pellet was resuspended in 1ml of MNase digestion buffer (0.32M sucrose, 50mM Tris-HCl, pH 7.5, 4mM MgCl_2_, 1mM CaCl_2_). The chromatin was digested by adding 1.5 μl micrococcal nuclease (NEB, M0247S) for 7 mins at 37 °C. The reaction was stopped by adding 10mM EDTA (pH 8.0), followed by centrifugation at 10,000 rpm for 10 min, retaining the supernatant (S1). Optionally, the pellet was suspended in 1ml Dialysis buffer (1mM Tris-HCl, pH 7.5, 0.2mM EDTA) and dialyzed against 1 Lt of Dialysis buffer overnight at 4°C. Samples were re-centrifuged at 10,000g for 10 min and the supernatant, S2, was optionally combined with S1. Mononucleosomes (S1 only) or mono-, di- and tri-nucleosomes (S1+S2) were purified on a 5–30% sucrose gradient containing 10 mM NaCl, 10 mM Tris-HCl (pH 7.4) and 0.2 mM EDTA and were used for immunoprecipitation. Mono-, di- and tri-nucleosomes extracts were diluted in 1:4 in N-ChIP Incubation Buffer (50mM NaCl, 50mM Tris-HCl, pH 7.5, 5mM EDTA) with 10μg of appropriate antibodies (custom-made anti-mH2A1.1, anti-mH2A1.2 and anti-mH2A2 antibodies (Ford *et al*, 2014; Valakos *et al*, 2023). Control experiments for the specificity of the antibodies were published in(Valakos *et al*, 2023)) and incubated with equilibrated protein-A dynabeads (Life technologies, 10001D) and competitors (BSA and yeast t-RNA) overnight at 4°C. Beads were collected on a magnet and washed two times each in Washing Buffer A (50mM Tris-HCl, pH 7.5, 10mM EDTA, 75mM NaCl), Washing Buffer B (50mM Tris-HCl, pH 7.5, 10mM EDTA, 125mM NaCl), and Washing Buffer C (50mM Tris-HCl, pH 7.5, 10mM EDTA, 175mM NaCl). Bound chromatin was eluted with freshly prepared elution buffer (1% SDS and 0.1M NaHCO_3_) at 65°C with vigorous shaking for 15 min. DNA was purified with QIAGEN Minelute PCR purification kit (QIAGEN, 28204) and then subjected to library preparation for Illumina sequencing (Ford *et al*, 2014). Briefly, 1-15 ng of ChIPed DNA was blunt-ended by End Repair Enzyme Mix [(DNA Polymerase, Klenow Fragment (NEB, M0210S), T4 DNA Polynucleotide kinase (NEB, M0201S)] followed by “A” tailing of 3’ ends and ligation with the annealed TruSeq Adapters (TruSeq ChIP Library Preparation Kit, Illumina, IP-202-1012). Conversion of the Y-shaped adapters to dsDNA occurred prior to the library size selection through 2.5% Metaphor (Lonza, 50180) /SeaKem LE (Lonza, BMA50004) (3:1 ratio) agarose gel electrophoresis. Each library was purified using QIAGEN miniElute columns (QIAGEN, 28204) and then subjected to pre-amplification. The final quantification of DNA libraries was carried out according to the Quantification Standards of Illumina on an Agilent Technologies 2100 Bioanalyzer (Agilent Technologies). Sequencing was performed in HiSeq 2000 (Illumina).

### Protein extraction and Western blot analysis

Cells were lysed in RIPA lysis buffer (150 mM NaCl, 50 mM Tris–HCl pH 8, 1% IGEPAL, 0.5% sodium deoxycholate, 0.1% SDS) that was added with a protease inhibitor cocktail (cOmplete Protease Inhibitor Cocktail, Roche, 11697498001). A total protein amount of 20 ug from each sample was denatured at 95°C for 10 min in Laemmli buffer containing beta-mercaptoethanol before electrophoresis. The primary antibodies anti-mH2A1.2 (Valakos *et al*, 2023) and anti b-tubulin (Cell Signaling Technology, 86298) (1:10,000) were diluted in PBST containing 2% skim milk and incubated at 4°C overnight. Commercial anti-mH2A1.2 antibody (Cell Signalling Technology, 4827) was used as a positive control. Secondary antibodies conjugated to horseradish peroxidase (HRP) (goat anti-rabbit HRP antibody, Biorad, 1706515; goat anti-mouse HRP antibody, Santa Cruz Biotechnology, sc-2055), and diluted in Tris-buffered saline with Tween 20 (TBST) (1:5,000) were incubated for 1h at RT. Signals were detected with Immobilon Western chemiluminescent HRP substrate (Millipore, WBKLS0500). B-tubulin was used as a loading control protein for normalization.

### Lentivirus production, infection and shRNA knockdown (KD)

KD datasets are derived from (Valakos *et al*, 2023). Briefly, scramble, anti-mH2A1.1 and anti-mH2A1.2 short hairpin (sh) RNA-producing DNA sequences were cloned in PLKO.1-puro-U6 plasmids. Reconstitution of lentiviruses was carried out in Human Embryonic Kidney 293T (HEK293T) cells by standard calcium phosphate DNA transfection protocols, using pMD2.G [a gift from Didier Trono (Addgene, 12259)] and psPAX2 [a gift from Didier Trono (Addgene, 12260)] packaging plasmids. Lentivirus-containing supernatant was harvested 3 days after medium change. Oligonucleotide sequences targeting mH2A1.1 or mH2A1.2 and scramble sequence are:

anti-mH2A1.1 shRNA Probe Sequence:

5’-TGACTTCTACACCGGTGGTGAACTCGAGTTCACCACCGGTGTAGAAGTCTTTTTG-3’

anti-mH2A1.2 shRNA Probe Sequence:

5’-GTGAGGTGGAGGCCATAATCAATCTCGAGATTGATTATGGCCTCCACCTCTTTTTG-3’

Scramble Probe Sequence:

5’-CCTAAGGTTAAGTCGCCCTCGCTCGAGCGAGGGCGACTTAACCTTAGGTTTTTG-3’

Control experiments validating the specificity of each oligonucleotide were published in (Valakos *et al*, 2023). mESCs were infected overnight with filtered viral supernatants and 8 ug/mL polybrene (Sigma-Aldrich, H9268), followed by medium change (for medium, see “Cell lines and mouse embryonic fibroblasts isolation” section). Three days post-infection, cells were selected with 5–10 μg/ml puromycin (Sigma-Aldrich, P8833) over a period of 7 days, followed by medium change and a two-day recovery period, after which they were harvested for downstream assays.

### Differentiation of mouse ESCs by Retinoic Acid

Differentiation of mouse ESCs was induced by retinoic acid (RA) (Santa-Cruz Biotechnology, sc-205589A), the main biologically active metabolite of vitamin A and one of the most important differentiation inducing factors (Rohwedel *et al*, 1999). To this aim, mouse ESCs (Bruce 4) were cultivated in the undifferentiated state ESC medium on a feeder layer of C57BL/6 MEFs treated with 10ug/mL Mitomycin C (Sigma-Aldrich, M4287) for 2.5–3 hours. Cells were trypsinized and 40,000 mouse ESCs were counted and plated on 0.1% gelatin-coated coverslips without feeders in a 24-well plate in ESC medium in the absence of LIF and in the presence of 10^-7^M RA. Medium was changed daily till day 6 when the differentiated phenotypes were analyzed with immunofluorescence. For immunofluorescence, cells were fixed with 4% paraformaldehyde for 15 min at RT, permeabilized with 0.3% Triton X-100, followed by three PBS washing steps (5 min each), blocked with 10% fetal bovine serum (FBS) for 1h at RT and finally labeled with Primary antibody against α-SMA (Abcam, ab7817; 1:400 dilution) in 1% FBS and 0.1% Triton X-100 overnight at 4°C. For detection of α-SMA, secondary antibody, Alexa Fluor 488-conjugated goat anti-mouse IgG H&L (Abcam, ab150113,1:500 dilution), in 1% FBS and 0.1% Triton X-100 was incubated for 1h at RT. Nuclei were stained with 1 mg/ml DAPI (4=,6-diamidino-2-phenylindole) (Merck, 268298)

### X-link Chromatin Immunoprecipitation (ChIP)

Naive MEFs and mESCs were cultured under optimal conditions and were fixed at a high-confluence stage at room temperature for 15 minutes using 1% formaldehyde, followed by quenching with 0.125M glycine at room temperature for 5 minutes. Upon extensive washes, cells were resuspended in Lysis [50mM Hepes (pH 7.9), 140mM NaCl, 1mM EDTA, 10% Glycerol, 0.5% NP-40, 0.25% Triton X-100] and then in sonication buffer [0.1% SDS, 1mM EDTA, 10mM Tris (pH 8.1)]. Chromatin shearing was carried out in the Covaris S220 sonicator using the Covaris TC12x12mm tubes (Tube AFA Fiber and Cap, Covaris, 520081) for 12 min (Duty Factor: 75, Peak: 25, Cycles per burst: 200) allowing the shearing of chromatin within a range of 250-500 base pairs DNA fragments. Triton X-100 and NaCl were then added in the sheared chromatin to final concentrations of 1% and 150 mM, respectively. ChIPs were carried out by incubating 100 μg of chromatin (corresponding to approximately 10x10^7^ cells) with 10μg of antibody per ChIP reaction overnight at 4°C, anti-E2F4 antibody (Cell Signaling Technology, 40291S) and custom-made rabbit IgG for isotype control. Next, Protein G-Dynabeads (Thermo Fisher Scientific, 10004D) pre-equilibrated in IP buffer [0.1% SDS, 1mM EDTA, 10mM Tris (pH 8.1), 1% Triton X-100, 150mM NaCl], were incubated with the chromatin-antibody solution in an orbital mixer at 4°C for 2 hours. The recovered resin was subsequently washed with low [0.1% SDS, 1% Triton X-100, 2 mM EDTA, 20 mM Hepes-KOH (pH 7.9), 150 mM NaCl] and high salt buffers [0.1% SDS, 1% Triton X-100, 2 mM EDTA, 20 mM Hepes-KOH (pH 7.9), 400 mM NaCl] and LiCl buffer [100 mM Tris-HCl (pH 7.5), 0.250 M LiCl, 1% NP-40, 1% Sodium Deoxycholate] and the captured chromatin fragments were subjected to Proteinase K (03115828001, Roche Life Sciences) digestion at 50°C for 15 minutes, followed by overnight incubation with RNase A (Macherey-Nagel, 740505) at 65°C. All DNA present in each sample was purified with Nucleomag beads (Macherey-Nagel, REF 744100.4) and eluted in TE buffer.

### ChIP-seq library construction

ChIP-seq library construction and sequencing for E2F4 ChIP-seq were carried out at the BRFAA Greek Genome Center (GGC). Libraries for E2F4 ChIP-seq were generated with NEBNext Ultra II DNA Library Prep Kit for Illumina, as indicated by the manufacturer’s guidelines (NEB, E7645L) using 10ng of DNA (chromatin that was not incubated with the antibody was used as input DNA). The quality of the libraries was assessed with the Agilent Bioanalyzer DNA 1000 assay (Agilent Technologies, 5067-1504) and quantitation was performed with the Qubit HS assay (Qubit 4.0 Fluorometer, Thermo Fisher Scientific, Q33238). Libraries were pooled in equimolar amounts for sequencing. At least 20 million, 75 bp long, Single-End reads were generated for each sample with illumina NextSeq 500 sequencer.

### ATAC-seq

ATAC-seq was performed as in (Klagkou *et al*, 2024). Briefly, cell pellets from Day 0, Day 1, Day 2, Day 3, Day 6, Day 9 and ESCs were lysed in Lysis Buffer [10 mM Tris-HCl (pH 7.5) 10 mM NaCl, 3 mM MgCl_2_, 0.1% NP-40, 0.1% Tween-20, 0.01% Digitonin (Promega, G9441)] for 3 min on ice, washed with Wash Buffer [10 mM Tris-HCl (pH 7.5) 10 mM NaCl, 3 mM MgCl_2_, 0.1% Tween-20] and centrifuged at 500 g for 10 min at 4 °C. Each pellet (nuclei) was resuspended in 50ul Transposition reaction mix [1x Tagment DNA Buffer (Illumina, 15027866), 16.5 uL 1x PBS, 0.1% Tween-20, 0.01% Digitonin (Promega, G9441),

2.5 uL Tn5 Transposase (Tagment DNA Enzyme 1, Illumina, 15027865)] and incubated at 37 °C for 30 min. DNA purification was performed using the Qiagen MinElute Reaction Cleanup Kit (Qiagen, 28204). Library purification was performed with the AMPure XP beads (Beckman Coulter, A63880). The quality of libraries was assessed using the Agilent Bioanalyzer DNA 1000 assay (Agilent Technologies, 5067-1504) and libraries were quantified with the Qubit™ High Sensitivity (HS) spectrophotometric method (Qubit 4.0 Fluorometer, Thermo Fisher Scientific, Q33238). Approximately 25 million, 100 bp long, Paired-End reads were generated for each sample with Illumina NovaSeq 6000 sequencer.

### ChIP-seq analysis

**mH2A ChIP-seq data**: The quality of the final fastq files was examined with the FASTQC software (v0.12.1). Fastq files were aligned against mm10 mouse genome with Bowtie2 aligner (Langmead & Salzberg, 2012) with the option “very-sensitive”. The output file was filtered for low quality reads, duplicates, blacklist regions and mitochondrial genome entries with the use of both from BEDtools (Quinlan & Hall, 2010) and SAMtools (Li *et al*, 2009). Peak calling was performed with MACS2 with default parameters (Zhang *et al*, 2008b) for both narrow peaks and broad peaks and the generated files were merged using the command bedtools merge. Bigwig files were generated with either deepTools2 either using IP/Input ratio normalization (Figs. 1B, E and EV3A, EV4A, EV3B) (Ramírez *et al*, 2016) or by subtracting the Input Signal from the respective IP signal using MACS2 bdgcmp function with the mode “subtract” (Zhang *et al*, 2008a) (Figs. 2B, 2E, 2G, 2H, 2I, 3D, 3F, 4D, 4F, 5C, 5D, 5E, EV2E, EV4F, EV4G, EV4H, EB4I, EV5D, EV5F). Heatmaps were created using deepTools2. Groups of regions with dynamic binding were defined by finding the exclusive peaks of each Day of reprogramming after intersection with those from Day 0. Groups A, B and C from mH2A1.2 data (Fig. 1E) were discovered through unsupervised k-means clustering. Overlap percentage with the OSKM peaks and Nanog at the ESCs state was found by using BEDtools intersect command. All peak files were filtered for duplicate coordinates and close regions with BEDtools mergeBed tool command (-d 146 option). mH2A1.2 repositioning genes were identified by first detecting common peaks between mH2A1.2 ChIP-seq profiles at Day 0 and in ESCs. The distances between peak summits at Day 0 and their corresponding ESC peaks were calculated, and summits separated by a distance over 10bps were classified as repositioned nucleosomes. Candidate regions were then manually validated through visual inspection of bigWig tracks in the Integrative Genomics Viewer (IGV). Functional Enrichment Analysis was performed with EnrichR (Chen *et al*, 2013a; Kuleshov *et al*, 2016) and the Dotplots were generated through R-studio ggplot2 package. Motif analysis was performed with HOMER motif discovery and analysis tool (v5.1) using default parameters and adding a redundancy filter of 0.4 (Heinz *et al*, 2010). Assignment of the mH2A sites to genes was performed with GREAT online tool (v4.0.4) “Single nearest gene” option for 3 kb maximum distance from TSS (McLean *et al*, 2010) unless stated differently. Functional Enrichment Analysis on target genes was performed with EnrichR (Chen *et al*, 2013a; Kuleshov *et al*, 2016). Annotation of peaks to genomic elements was performed with CEAS tool (Shin *et al*, 2009). Nucleosome occupancy analysis was performed using DANPOS3 (Chen *et al*, 2013b). mH2A1.1, mH2A1.2 and mH2A2 ChIP-seq datasets for MEFs and ESCs were used from (Valakos *et al*, 2023).

**OSKM, Nanog, E2F4 and NRF-1 ChIP-seq data**: For our O/S/K/M ChIP-seq analysis we utilized a combined dataset derived from multiple experiments either from our laboratory (Papathanasiou *et al*, 2021), or by others (Chen *et al*, 2016; Chronis *et al*, 2017; Di Giammartino *et al*, 2019; Knaupp *et al*, 2017), deposited under the GEO IDs GSE114581, GSE274131 GSE90893, GSE67520, GSE101905 and GSE113429. Analysis pipeline is presented thoroughly in (Klagkou *et al*, 2024), except for the Day 2 (Chronis *et al*, 2017) and Day 9 (Knaupp *et al*, 2017; Di Giammartino *et al*, 2019), which were analyzed *de novo* as in (Klagkou *et al*, 2024) for the current study. H3K27ac, H3K9ac, H3K4me1, and H3K4me2 ChIP-seq datasets were retrieved from (Chronis *et al*, 2017). NRF-1 ChIP-seq data were retrieved from (Baird *et al*, 2017; Domcke *et al*, 2015). Fastq files were aligned against mm10 mouse genome with Bowtie2 aligner (Langmead & Salzberg, 2012) with the option “very-sensitive”. The output file was filtered for low quality reads, duplicates, blacklist regions and mitochondrial genome entries with the use of both from BEDtools (Quinlan & Hall, 2010) and SAMtools (Li *et al*, 2009). Peak calling was performed with MACS2 with default parameters (Zhang *et al*, 2008b) for narrow peaks regarding OSKM, Nanog, E2F4 and NRF-1, and for broad peaks regarding H3K27ac, H3K9ac, H3K4me1, and H3K4me2. Bigwig files were constructed by subtracting the Input Signal from the respective IP signal using MACS2 bdgcmp function with the mode “subtract”.

### RNA-seq Analysis

RNA-seq for MEFs and time-points data were used from (Klagkou *et al*, 2024) (GSE274132), while ESCs data were used from (Valakos *et al*, 2023) (GSE215885). Quality Control was performed at the fastq raw data file for each sample using the “FASTQC” software. Fastq files were aligned to mm10 mouse genome using HISAT2 (Kim *et al*, 2015). Counts were defined using HTSeq htseq-count command with the “intersection non empty” option (Anders *et al*, 2015). The count files were used as Input for DESeq2 (Love *et al*, 2014). Normalization was performed with the “estimate size factor” function followed by Differentially Expressed Genes Analysis. Pathway analysis was performed at EnrichR (Chen *et al*, 2013a; Kuleshov *et al*, 2016). Heatmaps were constructed with pheatmap package in R after computing the respective z-score.

### ATAC-seq Analysis

ATAC-seq bioinformatics analyses were performed using The Galaxy Suite (Afgan *et al*, 2022). The quality of the sequencing reads was evaluated using the FastQC algorithm (Simon Andrews, 2020). Reads were trimmed to 50bp using the Trimmomatic tool (Bolger *et al*, 2014). Sequencing reads were mapped to the mm10 version of the mouse genome using the Bowtie2 algorithm with the “very sensitive end to end” option (Langmead & Salzberg, 2012). The Samtools Fixmate command was used prior to duplicate removal with the MarkDuplicates command from Picard tools using the option “do not write duplicates to the output file” (Li *et al*, 2009). Reads from all samples were downsampled to the same sequencing depth using the Downsample BAM option from Picard tools. Peaks were called using the MACS2 algorithm with the options “no model”, “extension-200”, “shift-100” and a q-value cut-off of 0.05. Bigwig files were constructed using the bamCoverage command from the Deeptools2 suite with RPKM as the normalization method.

## Data Availability

All reagents generated in this study are available upon request. The new NGS data produced and presented in the study (raw and analyzed files) are deposited in the Gene Expression Omnibus data repository under the GEO accession ID GSE296734. Already published data used in the study (either analyzed again, or used as published) are available in the Gene Expression Omnibus data repository under the GEO accession IDs GSE274130, GSE89344, GSE67867, GSE114581, GSE274131, GSE274132 GSE90893, GSE67520, GSE101905, GSE113429, GSE215885. Token for reviewer access of GSE296734: wxaneqgkhhupbut

## Author Contributions

Study conception and design: D.T., D.V. and A.K.; Performed experiments and data collection: D.V., A.K., E.K., and G.V.; Multi-omics & statistical analyses: D.V., and A.P.; Writing-original draft: D.V. and D.T.; Writing-review and editing: all authors; Funding acquisition: D.T; Supervision, D.T.

## Disclosure and competing interest statement

The authors declare no competing interests.

## Acknowledgments

We thank Joseph Papamatheakis, Charalampos Spilianakis, George Mosialos, Spyros Foutadakis, Maria Pliatska, Ioanna Polidouri and Antonia Katsouda for critical reading of the manuscript. We also thank Anastasios Delis and Stamatis Pagakis for consultation regarding Confocal Microscopy. This work was supported by grants to D.T. from the Greek General Secretariat for Research and Innovation (GSRI) (Cooperative Grants Synergasia I #969, Emblematic Action Against COVID19, BIOIMAGING.GR (MIS-5002755)), Human Foundation for Research and Innovation (015999), European Committee FP7 projects (STPredicta, and Biofos), from the European Economic Area (EL0084), GSRI Public Investments Program SAM 013 and from the KMW offsets program. E.K. was also supported from the State Scholarships Foundation (Operational Programme MIS-5000432). D.V. was also supported by a scholarship from the Bodossaki Foundation.

## Expanded View Figure Legends

**Figure EV1:**
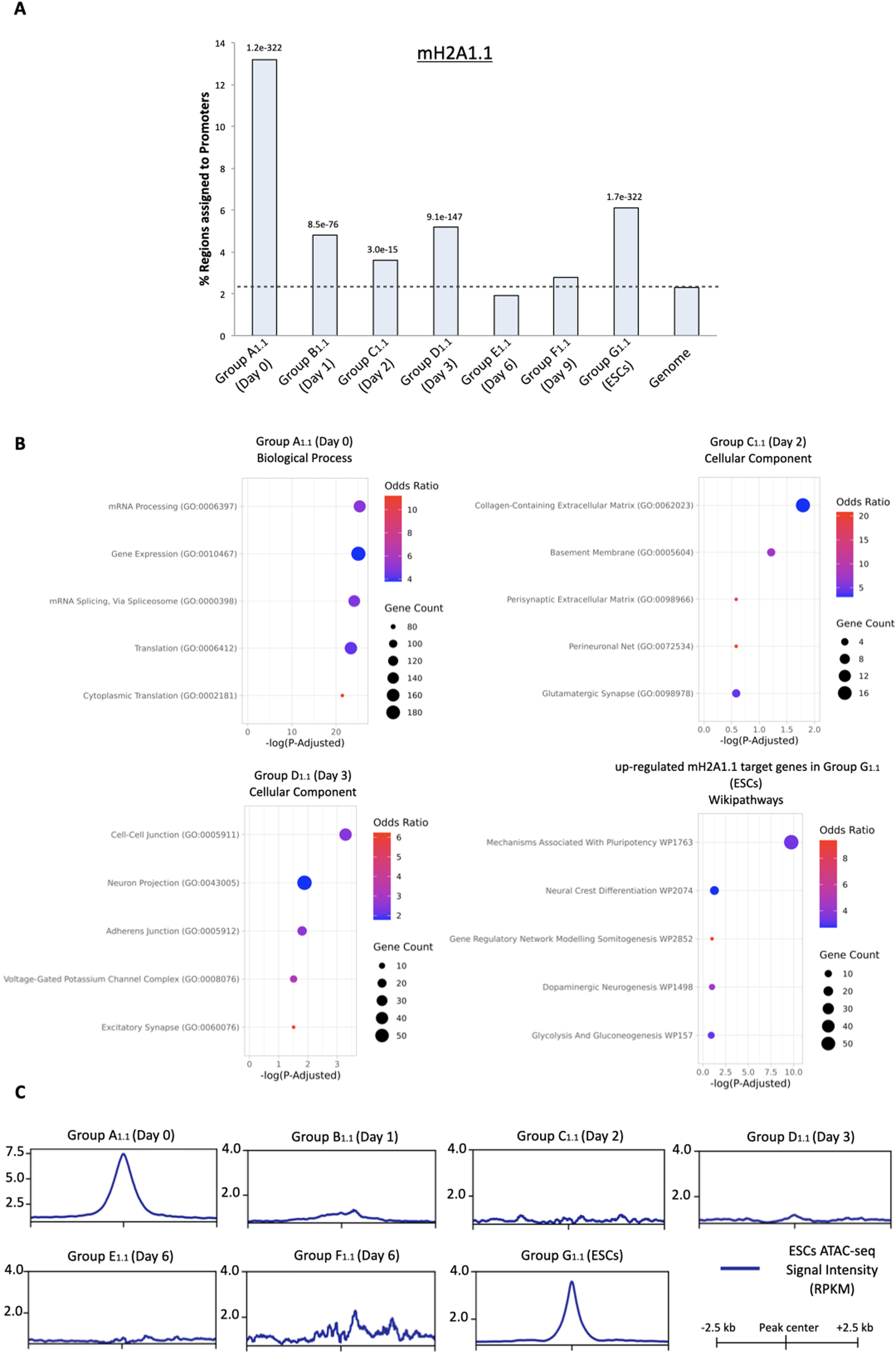
Complementary analysis of mH2A1.1 regions as defined in Fig. 1B (Relevant to Figure 1) (A) Bar graphs indicating the percentage of promoter regions within Groups A_1.1_–G_1.1_ (as defined in Fig. 1B). The last bar (“Genome”) represents the genomic baseline of promoter frequency in the mouse genome; comparison to this baseline (dashed line) highlights enrichment in each group. P-value as defined from CEAS tool is depicted on the top of bars representing regions with significant enrichment in promoter elements and used for downstream analyses. (B) Functional enrichment analysis (Over-Representation Analysis, ORA) for genes located near mH2A1.1 nucleosome sites in each group. Shown are the top five significantly enriched terms per group, ranked by P-Adjusted value (Cut-off for statistical significance: P-Adjusted < 0.05). (C) Summary plots depicting the ATAC-seq signal (RPKM) regions found in ESCs across Groups A_1.1_–G_1.1_ regions (as defined in Figure 1B), centered at mH2A1.1 peak summits and spanning –2.5 kb to +2.5 kb.

**Figure EV2:**
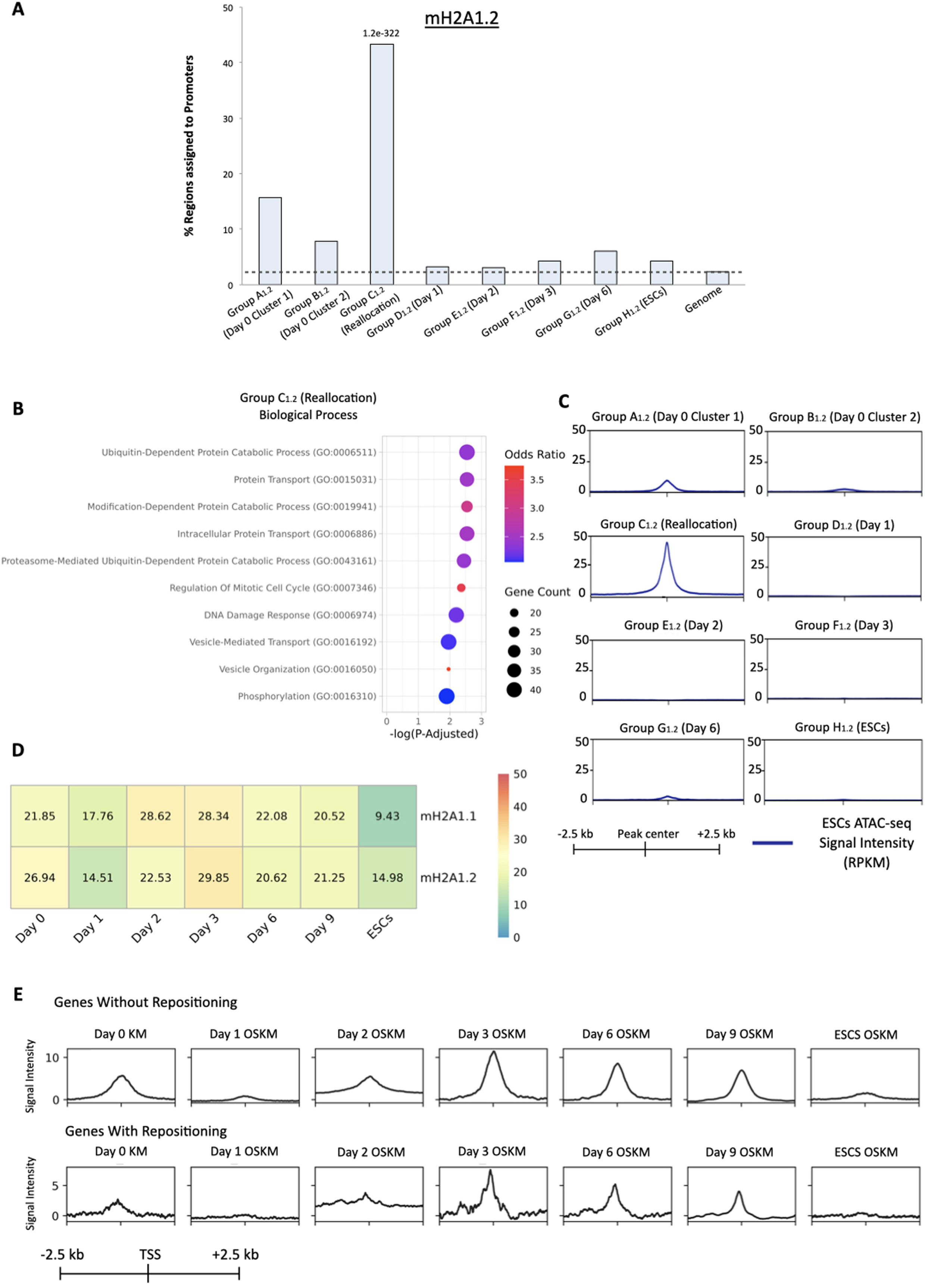
Complementary analysis of mH2A1.2 regions as defined in Fig. 1E (Relevant to Figure 1) (A) Bar graphs indicating the percentage of promoter regions within Groups A_1.1_–H_1.1_ (as defined in Fig. 1E). The last bar (“Genome”) represents the genomic baseline of promoter frequency in the mouse genome; comparison to this baseline (dashed line) highlights enrichment in each group. P-value as defined from CEAS tool is depicted on the top of bars representing regions with significant enrichment in promoter elements and used for downstream analyses. (B) Functional enrichment analysis (ORA) for genes associated with Group C_1.2_ (relocation) regions. Shown are the top 10 significantly enriched biological processes, ranked by P-Adjusted (Cut-off for statistical significance: P-Adjusted < 0.05). (C) Line plots showing ATAC-seq signal (RPKM) in ESCs across Group A_1.2_-H_1.2_ regions (as in Fig. 1E), centered at mH2A1.2 peaks and spanning –2.5 kb to +2.5 kb. (D) Heatmap depicting the percentage of mH2A1.1 peaks which are common with the mH2A1.2 peaks (upper row) and the percentage of mH2A1.2 peaks which are common with the mH2A1.1 peaks (lower row), at Day 0, during reprogramming stages (Day 1, Day 2, Day 3, Day 6, Day 9) and ESCs. (E) Summary plots of OSKM ChIP-seq signal at the TSS of genes without repositioning (upper panel) and genes with repositioning (lower panel). ChIP-seq signal was normalized by subtracting the Input signal of the respective time-point from the IP signal.

**Figure EV3:**
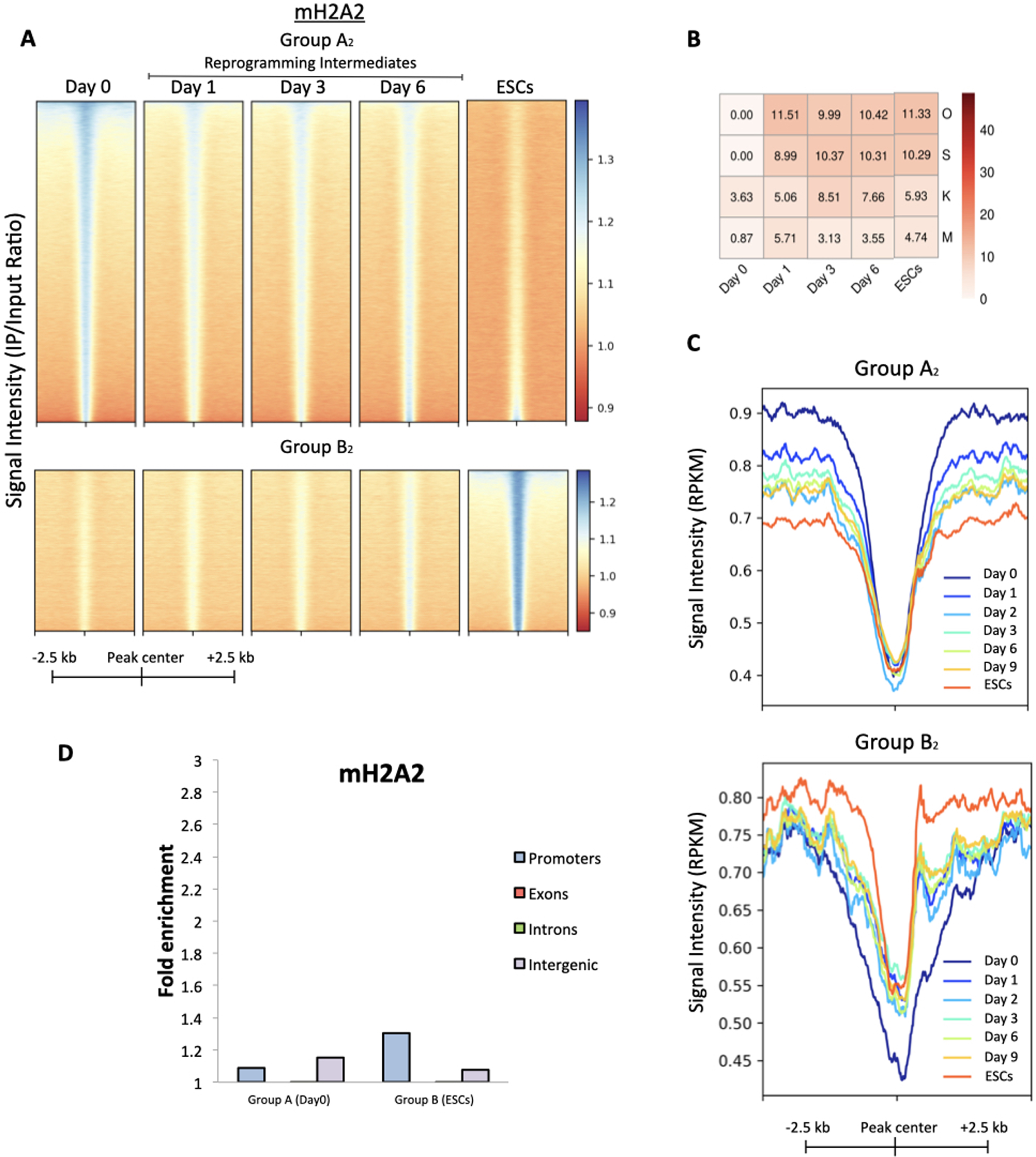
Analysis of mH2A2 regions (Relevant to Figure 1) (A) Heatmaps illustrating the genome-wide redistribution of mH2A2-containing nucleosomes across different genomic regions (Group A_2_ and Group B_2_) during reprogramming. Regions are centered on mH2A2 peaks and span –2.5 kb to +2.5 kb from the peak center. Binding sites were grouped based on occupancy dynamics, sorted in descending order of ChIP-seq signal intensity (IP/Input) in MEFs and maintaining the same order for all time-points and peaks were identified using MACS2. (B) Heatmap showing the percentage of shared binding peaks between mH2A2 and each of the OSKM factors (Oct4, Sox2, Klf4 and c-Myc) throughout reprogramming (Day 0, Day 1, Day 2, Day 3, Day 6, Day 9 and ESCs). OSKM factors are indicated on the right. (C) Summary plots of ATAC-seq signal (RPKM) across Group A_2_ (MEF-specific) and Group B_2_ (ESC-specific) mH2A2 regions as defined in (A). Chromatin accessibility is shown for MEFs (Day 0), Day 1, Day 2, Day 3, Day 6, Day 9 and ESCs across a ±2.5 kb window centered on the mH2A2 peak summits. (D) Bar graph showing the fold enrichment in genomic annotations of mH2A2 Groups A_2_–B_2_ regions (as defined in Figure EV3A), categorized into promoters, exons, introns and intergenic regions.

**Figure EV4:**
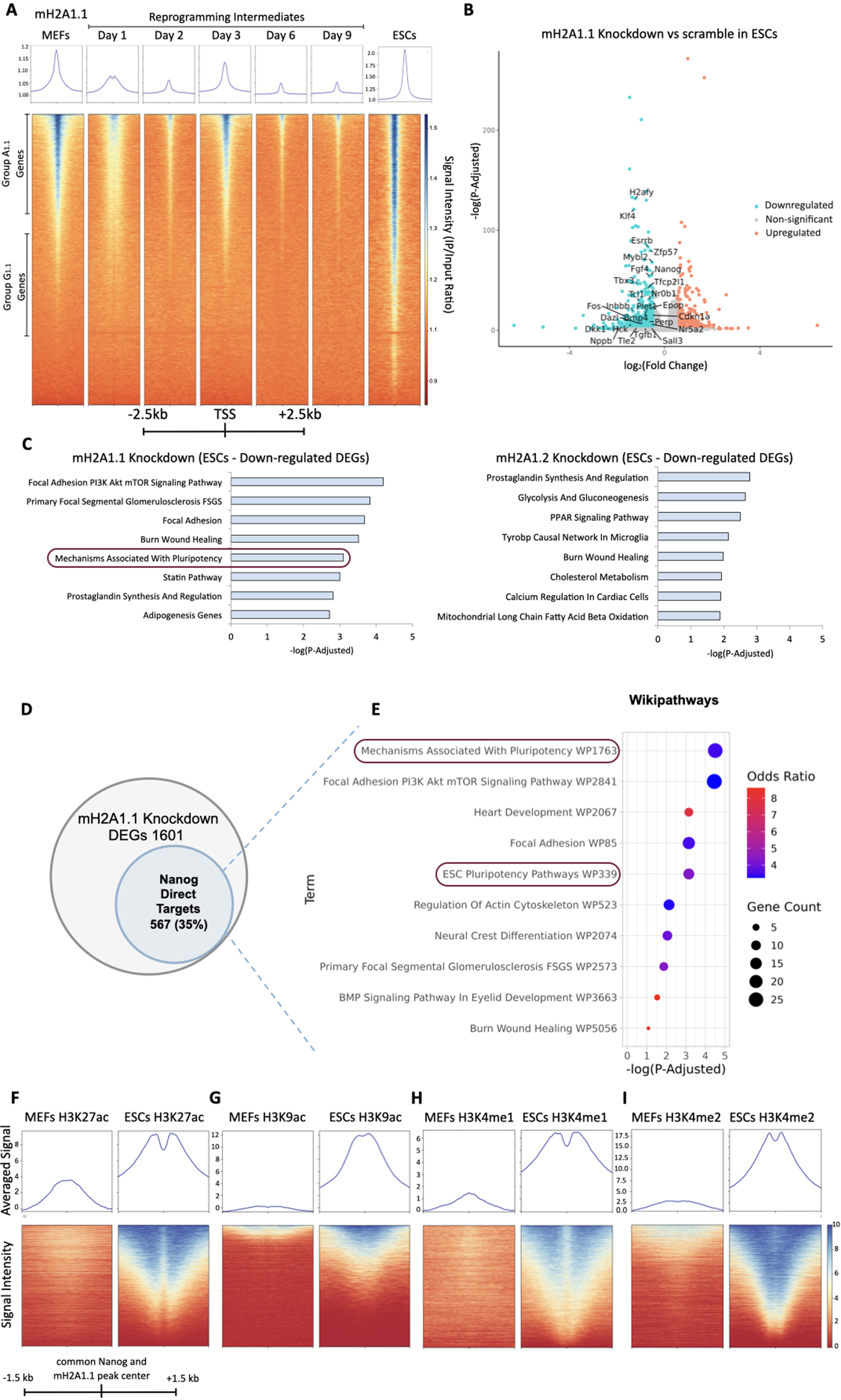
Composite Nanog/mH2A1.1 nucleosome elements protect pluripotency (Relevant to Figure 2) (A) Summary plots and heatmaps displaying mH2A1.1 ChIP-seq signal (IP/Input ratio) across all annotated mouse promoters from –2.5 kb to +2.5 kb relative to the TSS of all genes throughout reprogramming (Day 0, Day 1, Day 3, Day 6, Day 9 and ESCs). Regions were sorted in descending order based on IP signal at MEFs state, with the same order maintained across time-points. Signal intensities are shown as IP/Input ratio. (B) Volcano plot displaying differentially expressed genes (DEGs) in mH2A1.1 Knockdown versus scramble control in ESCs. Pluripotency-associated DEGs highlighted in Figure EV4E ("Mechanisms associated with Pluripotency") are indicated (purple box). Cut-offs: P-Adjusted < 0.05, log_2_FC > 0.5 or < –0.5. (C) Functional enrichment (Wikipathways) of downregulated genes in mH2A1.1 KD ESCs (left panel) compared to mH2A1.2 KD ESCs (right panel). Top 8 terms are shown, sorted by adjusted p-value (P-Adjusted < 0.05). (D) Venn Diagram depicting the number of DEGs derived from mH2A1.1 KD in ESCs and their percentage as direct Nanog target genes. Target genes were defined from Nanog ChIP-seq in ESCs using GREAT. Genes where Nanog peaks were identified within 5kb from the TSSs were considered as direct targets. (E) Functional Enrichment Analysis (Over-representation Analysis, ORA) of DEGs derived from mH2A1.1 KD in ESCs and are direct targets of Nanog. (F) Summary plots and heatmaps displaying H3K27ac ChIP-seq signals from -1.5kb to +1.5 kb relative to the center of the common Nanog and mH2A1.1 peaks in MEFs (left panels) and ESCs (right panels). ChIP-seq signal was normalized by subtracting Input from IP. (G) Same as (F), but for H3K9ac ChIP-seq signal. (H) Same as (F), but for H3K4me1 ChIP-seq signal. (I) Same as (F), but for H3K4me2 ChIP-seq signal. H3K27ac, H3K9ac, H3K4me1 and H3K4me2 ChIP-seq datasets were retrieved from(Chronis *et al*, 2017).

**Figure EV5:**
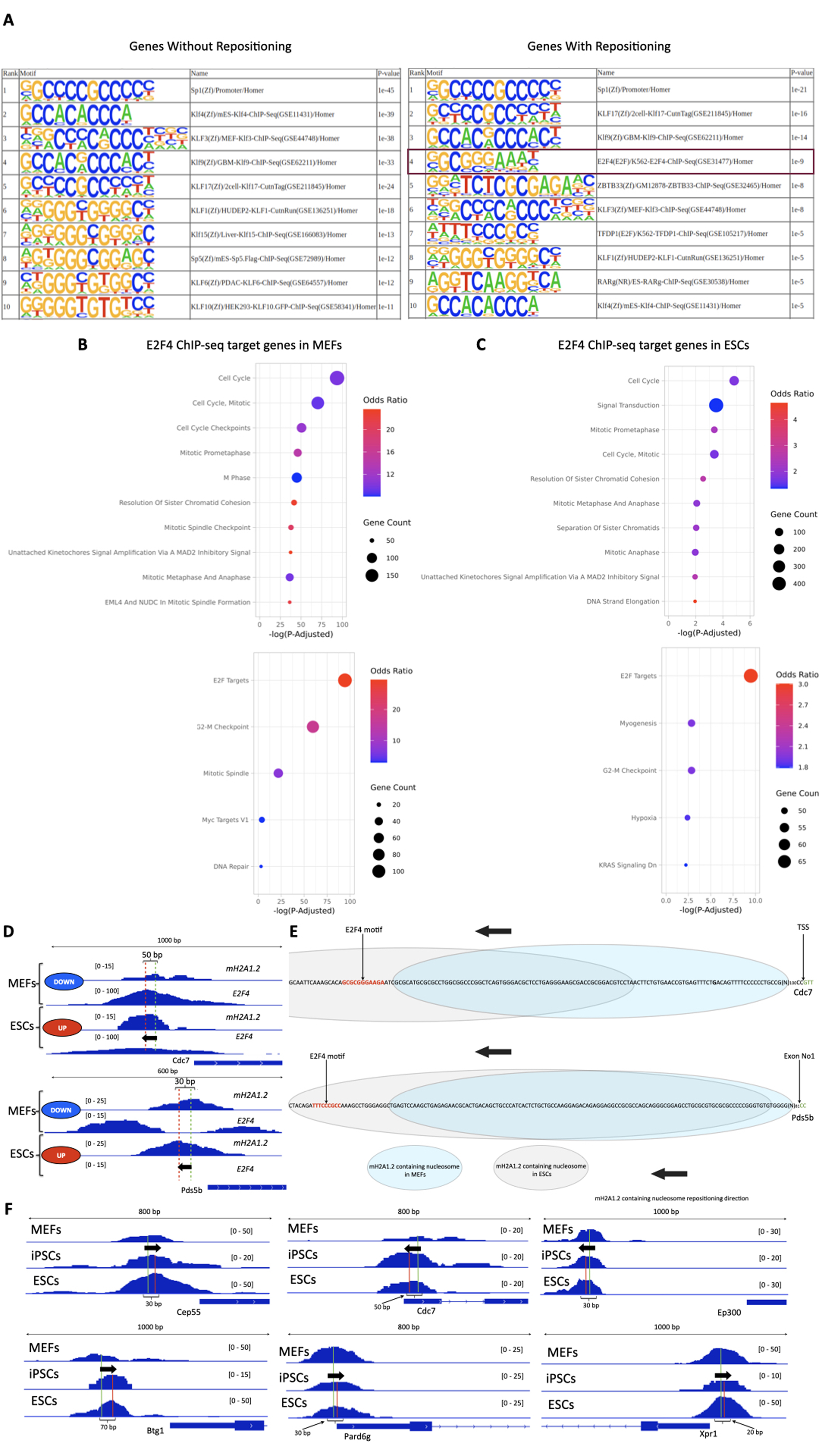
mH2A1.2 nucleosome repositioning switches on or off E2F4 DNA binding during reprogramming (Relevant to Figure 4) (A) Comparative motif analysis of DNA sequences at mH2A1.2-bound nucleosomes in genes without repositioning (left) and those with mH2A1.2 nucleosome repositioning (right). E2F4 motifs are specifically enriched in the repositioned group (highlighted in bold purple box), suggesting a role in transcriptional regulation. (B) Functional enrichment analysis of E2F4 target genes in MEFs using the Reactome (top) and MSigDB (bottom) databases. Enriched pathways are related to cell-cycle regulation and repression. (C) Functional enrichment analysis of E2F4 target genes in ESCs using the Reactome (top) and MSigDB (bottom) databases. Enriched pathways indicate transcriptional activation of cell-cycle progression genes. (D) IGV browser snapshots of representative genes showing mH2A1.2 nucleosome repositioning and corresponding changes in E2F4 DNA binding between MEFs and ESCs. All tracks were normalized by subtracting Input from the IP signal. Black arrows indicate the direction of nucleosome repositioning; green and red dashed lines mark original and final peak centers, respectively. Repositioning distances were calculated based on mH2A1.2 ChIP-seq peak centers. In ESCs, nucleosome repositioning inhibits E2F4 binding, enabling gene activation. (E) Schematic illustration of mH2A1.2 nucleosome repositioning at selected loci from (D). In MEFs, mH2A1.2 nucleosomes (pale blue ovals) do not mask E2F4 motifs (red sequence), allowing E2F4 binding. In ESCs, repositioned mH2A1.2 nucleosomes (gray ovals) occlude the motifs, blocking E2F4 access. (F) Comparative IGV browser views showing mH2A1.2 nucleosome repositioning from MEFs to iPSCs and ESCs. Arrows indicate the repositioning direction, and dashed lines represent original (green) and final (red) peak centers. Repositioning distances were calculated from ChIP-seq peak shifts between cell states.

## Expanded View Datasets

### Dataset EV1

The different Groups of mH2A1.1 peaks as presented in Fig. 1B.

### Dataset EV2

The different Groups of mH2A1.2 peaks as presented in Fig. 1E.

### Dataset EV3

The different Groups of mH2A2 peaks as presented in Fig. EV3A.

### Dataset EV4

List of genes without mH2A1.2 nucleosome repositioning (left) and with mH2A1.2 nucleosome repositioning (right). NRF-1 binding status is also indicated in a separate column.

